# Calcitonin-Driven Reactivation of the Hippo Tumor-Suppressor Cascade Attenuates YAP/TAZ Oncogenic Signaling in Glioblastoma

**DOI:** 10.1101/2025.11.20.689524

**Authors:** Jayita Goswami, Prithviraj Uttarasili, Lakshay Garg, Kaval Reddy Prasasvi, Abhishek Chowdhury, Subhrodeep Saha, Jayanta Chatterjee, Anand Srivastava, Kumaravel Somasundaram

## Abstract

Activation of WT calcitonin receptor (WT CTR) by Calcitonin (CT) acts as growth inhibitory axis in GBM, while the patient-derived loss-of-function CTR mutants promote tumor growth and lower patient survival. The precise mechanisms that orchestrate CTR activation, modulate its downstream signaling dynamics, and give rise to structural anomalies in mutant variants remain incompletely defined and warrant comprehensive investigation. Here, we found that salmon CT treatment suppressed the growth of patient-derived glioma stem cells (GSCs) by activating the Hippo pathway through the CTR/cAMP/PKA/LATS1 signaling cascade, while simultaneously inhibiting the oncogenic YAP/TAZ transcription factors. However, GSCs expressing a phosphorylation-resistant YAP mutant were refractory to growth inhibition by CT in vitro. GSCs, unlike differentiated glioma cells (DGCs), expressed high levels of CTR thus making them suitable for CT-based therapy. Furthermore, the intranasal delivery of salmon CT inhibited glioma growth initiated by GSCs in an intracranial orthotopic mouse model, resulting in increased survival in the mice. In addition, unlike the WT, the CTR mutants failed to activate the Hippo pathway. A large-scale, microsecond-long all-atom molecular dynamics simulation study of mutant CTR systems revealed remarkable changes in CTR interactions with the CT and Gα-GTPase domains. Packing and conformational dynamics data from simulation studies of the WT and mutant CTR systems could explain the perturbed cAMP/PKA signaling that may compromise downstream signaling. Together, we demonstrate that the CT/CTR axis inhibits glioma by activating the Hippo pathway, provide evidence for the structural defects in the loss-of-function CTR mutants, and propose a CT-based therapeutic strategy for GBM.

**Statement of significance:** CT/CTR axis targets YAP/TAZ through Hippo pathway activation. Patient-derived GSCs express higher CTR levels, and intranasally delivered CT suppresses GSC-initiated tumors. All-atom molecular dynamics simulation uncovers structural perturbations in the CTR mutants.

## Introduction

G protein-coupled receptors (GPCRs), which represent the largest protein family, have been implicated in several diseases, such as cancer, and others **[1]**. GPCRs have been considered targets for developing drugs as they control a wide range of physiological processes. While 34% of FDA-approved drugs target GPCRs, only eight of them have been developed as anticancer drugs **[2]**. Aberrant gene expression due to copy number aberrations, epigenetic mechanisms, and genetic mutations in GPCRs has been shown to induce ligand-dependent or independent constitutive signaling, contributing to oncogenesis **[2–6]**.

Calcitonin receptor (CTR), a member of the class B family of GPCRs, binds calcitonin (CT), a neuroactive peptide, to maintain calcium homeostasis in bone and kidney **[7]**. While the CTR expression has been demonstrated in many cancers, the role of CTR in cancer initiation and development remains to be established. While CT treatment has been shown to inhibit apoptosis and promote prostate tumor growth **[8]**, the CT/CTR axis inhibited the growth of breast cancer, glioma, and Giant cell tumors **[9–11]**. Loss-of-function mutations in Glioblastoma (GBM) identified a subgroup with a poor prognosis **[10]**. The glioma cell growth inhibition by CT treatment in a WT CTR-dependent manner is correlated with inhibition of JNK, ERK, and AKT phosphorylation **[10]**. WT CTR, but not CTR mutants, inhibited the Ras-mediated transformation of immortalized astrocytes **[10]**. However, the precise mechanism of growth inhibition by CT and signaling downstream of CTR is yet to be elucidated.

TCGA database-based analysis revealed that 20% of all sequenced human tumors carry mutations in GPCR genes **[2]**. A survival correlation analysis of mutated genes and pathways identified that patients with a defective “neuroactive ligand interaction pathway” had a worse prognosis **[10]**. A focused investigation revealed that CTR with patient-derived mutations failed to respond to CT and inhibit glioma growth **[10]**. Recently, a near-atomic-resolution structure of CT-bound CTR was obtained **[12]**. CT binding to extracellular loops creates structural alterations of the transmembrane domain, facilitating the interaction between Gα and intracellular loops of CTR **[12–14]**. However, the impact of loss-of-function mutations on the CTR structure is unclear.

In this study, we found that the CT/CTR axis inhibits oncogenic transcriptional coactivators, YAP/TAZ, by activating the Hippo pathway through the cAMP/PKA/LATS1 signaling pathway. Intranasal delivery of salmon CT inhibited the growth of human and murine Glioma stem cells (GSCs)-initiated tumors in an intracranial mouse model. Large-scale all-atom molecular dynamics (AAMD) simulation studies revealed that each patient-derived mutant noticeably changes the CTR packing and its conformational dynamics. Depending on the location of the mutations in the CTR, the mutant system either compromises CT binding, Gα binding, or the strength of membrane lipid interactions.

## Methods

### Experimental model and subject details

The animal studies were conducted on 6–8-week-old female C57BL/6J mice and Athymic Nude mice (NIH nu/nu), with approval from the Institute’s Ethical Committee for Animal Experimentation under Project Number CAF/Ethics/743/2020. The animals were kept under a 12-hour light-dark cycle and provided unlimited access to a standard diet. All the experiments were conducted during the light phase of the cycle.

### Plasmids

8xGTIIC-luciferase (TEAD promoter Luc plasmid), mEGFP-N1, mEGFP-N1-YAP, and mEGFP-N1-YAPS127A plasmids were bought from Addgene, USA (#43806). The CTGF Promoter Luc construct was created in our lab **[15]**.

### Other methods and materials

The details of other methods and materials used, as described below, are given in the supplementary section: Cell lines, RPPA analysis, Neurosphere culturing, GSC Differentiation/dedifferentiation, Limiting Dilution Assay, Lentivirus preparation and transduction of cells, Luciferase and beta-galactosidase (β-gal) assay, RNA isolation, cDNA conversion, and Real-qPCR and the list of primers, Western Blotting, Apoptosis assay, Colony Formation assay, Cell Viability assay, Immunofluorescence, Intracranial injection, In vivo Imaging, Haematoxylin and Eosin staining, Survival Analysis, Quantification and Statistical Analysis, Molecular Modelling of CTR, Insilico reconstitution of full-length CTR from structurally solved fragments, In silico reconstitution of mutant CTR with binding partners, and molecular dynamics simulation details

## Results

### CT/CTR signaling activates the Hippo pathway to inhibit YAP/TAZ and suppresses glioma growth

We previously demonstrated that the CT/CTR signaling axis exerts a growth-suppressive effect in glioma [**10**]. To elucidate the downstream molecular events and effector proteins mediating this response, we subjected extracts of CT-treated LN229 glioma cells to Reverse Phase Protein Array (RPPA) profiling. This unbiased analysis identified a spectrum of regulated proteins that were subjected to Gene Ontology (GO). GO analysis revealed a significant enrichment of the **Hippo signaling pathway** (**Figure 1A; Supplementary Table 1**), suggesting that the CT/CTR axis modulates Hippo pathway activity. The Hippo pathway is an evolutionarily conserved signaling cascade that governs organ size and tissue homeostasis, and its deregulation is implicated in many diseases, including cancer [**15–17**]. Mechanistically, pathway activation involves the MST1/2–LATS1/2 kinase cascade-dependent phosphorylation of YAP/TAZ proteins, which subsequently undergoes cytoplasmic retention by 14-3-3 proteins, followed by ubiquitin-mediated proteasomal degradation [**18**].

**Figure 1.**
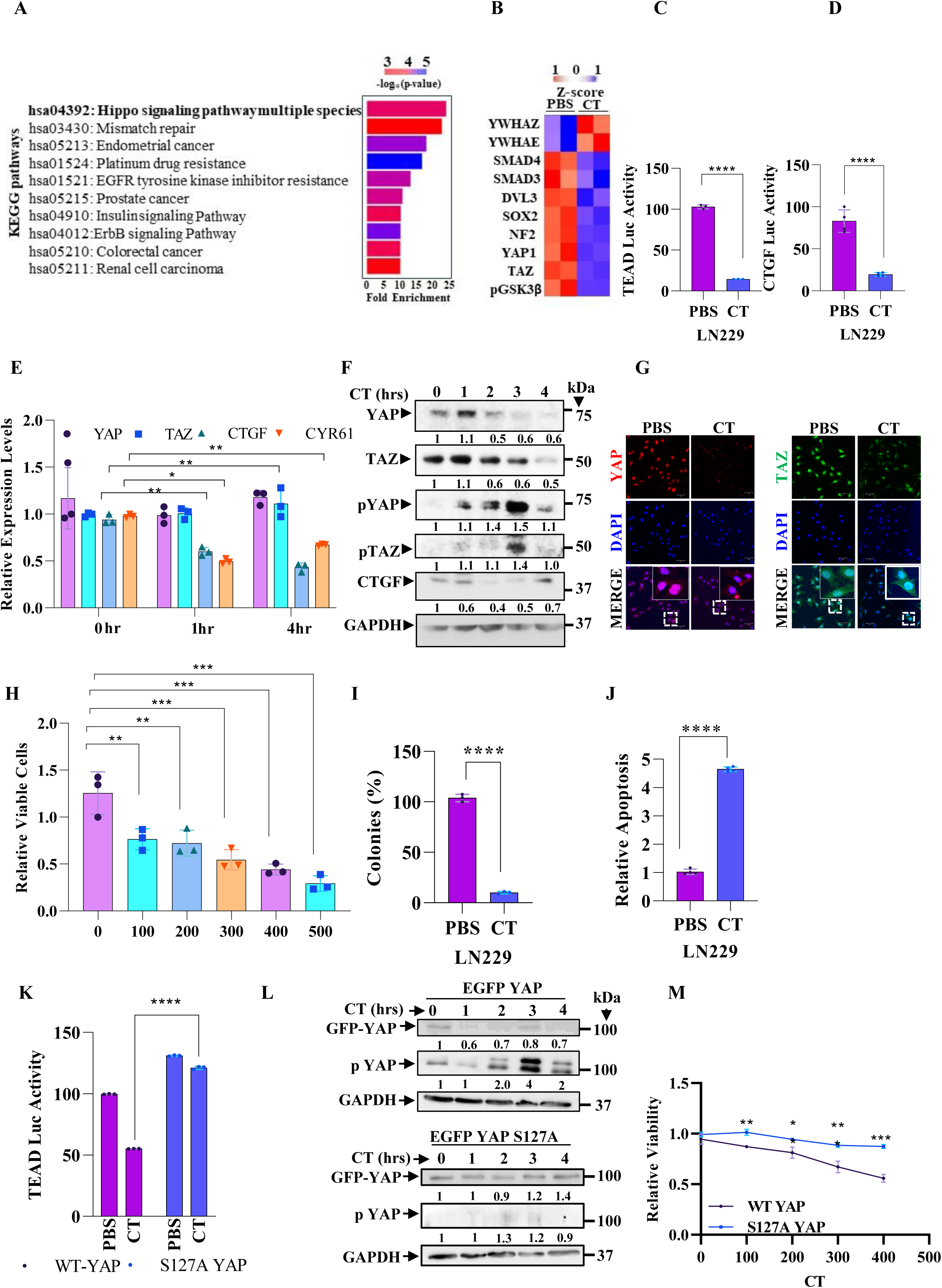
CT inhibits YAP and TAZ in a CTR-dependent manner. **(A)** Differentially expressed proteins identified through Reverse Phase Protein Array (RPPA) analysis in CT-treated LN229 cells and subsequently analysed using DAVID for KEGG pathway enrichment reveal significant enrichment (p<0.05) of the Hippo pathway following CT treatment. **(B)** Heatmap showing differentially regulated (|log_2_FC| ≥0.58, p<0.05) genes of the Hippo pathway after CT treatment, assessed from RPPA data. Red and blue represent upregulated and downregulated genes under the CT-treated condition. Effect of PBS or CT on GBM cell line LN229 transfected with TEAD Luc (t test; n=3/group) (**C**) and CTGF Luc (**D**) construct (t test; n=4/group). (**E)** RT-qPCR analysis shows YAP, TAZ, CTGF, and CYR61 transcript levels upon CT treatment (300nM) in the LN229 cell line for the indicated time points (two-way ANOVA n=3/group). (**F)** Western blotting shows YAP, TAZ, pYAP, pTAZ, and CTGF upon CT treatment for the indicated time points. Confocal microscopy analysis shows YAP and TAZ **(G)** levels in LN229 cells with and without CT treatment. Red indicates YAP, green indicates TAZ, and blue indicates DAPI. The merged images are shown for representation. Magnification x20, scale bar 20 µm. (**H)** MTT assay showing the viability of LN229 cells at the indicated concentrations of CT after 72 hours of treatment (one-way ANOVA n=4/group). (**I)** Quantification of colony formation under PBS and CT-treated conditions (t test; n=4/group). (**J)** Quantification of apoptotic cells by annexin V and PI staining after CT treatment for 72 hours (t test; n=4/group). **(K)** The plot shows TEAD Luc activity from LN229 cells co-expressing WT YAP or constitutively active YAP S127A (two-way ANOVA; n=3/group). **(L)** Western blot showing the effect of CT on LN229 cells expressing WT or S127A YAP. **(M)** The graph shows the effect of CT on the viability of LN229 cells expressing WT or S127A YAP (two-way ANOVA; n=3/group).

RPPA data analysis demonstrated a marked reduction in YAP, TAZ, and SOX2 (an established YAP target), concomitant with increased expression of YWHAZ and YWHAE, two members of the 14-3-3 protein family in CT-treated cells, which supported the presence of an activated Hippo pathway (**Figure 1B; Supplementary Table 2**). Hippo pathway activation was confirmed by significant decrease in TEAD-Luc and CTGF-Luc promoter (YAP/TAZ-dependant reporters) activity (**Figure 1C, D**; **Supplementary Figure 1A, B**), downregulation of the canonical YAP/TAZ target genes, CTGF and CYR61 with no change in YAP and TAZ transcript levels (**Figure 1E; Supplementary Figure 1C**), decreased abundance of total YAP and TAZ with a simultaneous enhanced phosphorylation of both proteins (**Figure 1F; Supplementary Figure 1D**, diminished nuclear localization of YAP (**Figure 1G; Supplementary Figure 1E**), and reduced CTGF protein levels (**Figure 1F; Supplementary Figure 1D**) in CT-treated LN229 and T98G glioma cell lines.

Next, we found that CT treatment significantly suppressed proliferation, inhibited colony-forming ability, and induced apoptosis (**Figure 1H, I, and J; Supplementary Figure 1F**) but not in CTR-silenced cells (**Supplementary Figure 1G, H, and I),** establishing the requirement of CTR for the growth-inhibitory activity of CT. Further, CTR knockdown abrogated CT-induced Hippo pathway activation, as evidenced by the inefficient suppression of YAP/TAZ-dependent reporter activity (**Supplementary Figure 1J**) and the absence of reductions in total YAP/TAZ or increases in pYAP/pTAZ protein levels (**Supplementary Figure 1K**). To rigorously confirm the requirement of YAP/TAZ targeting, we used a phosphorylation-resistant YAP mutant (S127A), which constitutively drives transcriptional activation and is refractory to Hippo pathway–mediated inhibition [**19**]. In cells expressing the S127A YAP mutant, CT treatment failed to reduce TEAD-Luc reporter activity, diminish total YAP, enhance pYAP, or impair proliferation (**Figure 1K, L, and M**). Collectively, these findings establish that the CT/CTR axis executes a tumor-suppressive effect by the Hippo signaling activation and the subsequent YAP/TAZ targeting

### Hippo pathway activation by CT/CTR involves cAMP/PKA/LATS1 signaling

To elucidate the molecular basis of Hippo pathway activation by CT, we examined the phosphorylation status of the upstream kinases MST1 and LATS1. CT treatment significantly increased phosphorylated LATS1 (pLATS1) levels but not the total LATS1 protein level (**Figure 2A**) with no change in phosphorylated and total MST1, indicating that CT activates the Hippo pathway at the LATS1 level and is MST1-independent. In glioma cells with LATS1 knockdown, CT treatment failed to suppress TEAD-luciferase reporter activity, decrease total YAP protein, reduce nuclear YAP localization, or inhibit proliferation (**Figure 2B, C, and D; Supplementary Figure 2A, B**). Complementary pharmacological inhibition of LATS1/2 with TRULI similarly abolished the effects of CT, including repression of TEAD reporter activity, downregulation of YAP/TAZ, increase in pYAP and pTAZ levels, exclusion of YAP from the nucleus, and inhibition of cell growth (**Supplementary Figure 2C, D, E, and F**). These findings establish LATS1 as a key mediator of CT/CTR-induced Hippo pathway activation, underscoring its essential role in transducing the anti-tumor effects of CT signaling in glioma cells.

**Figure 2.**
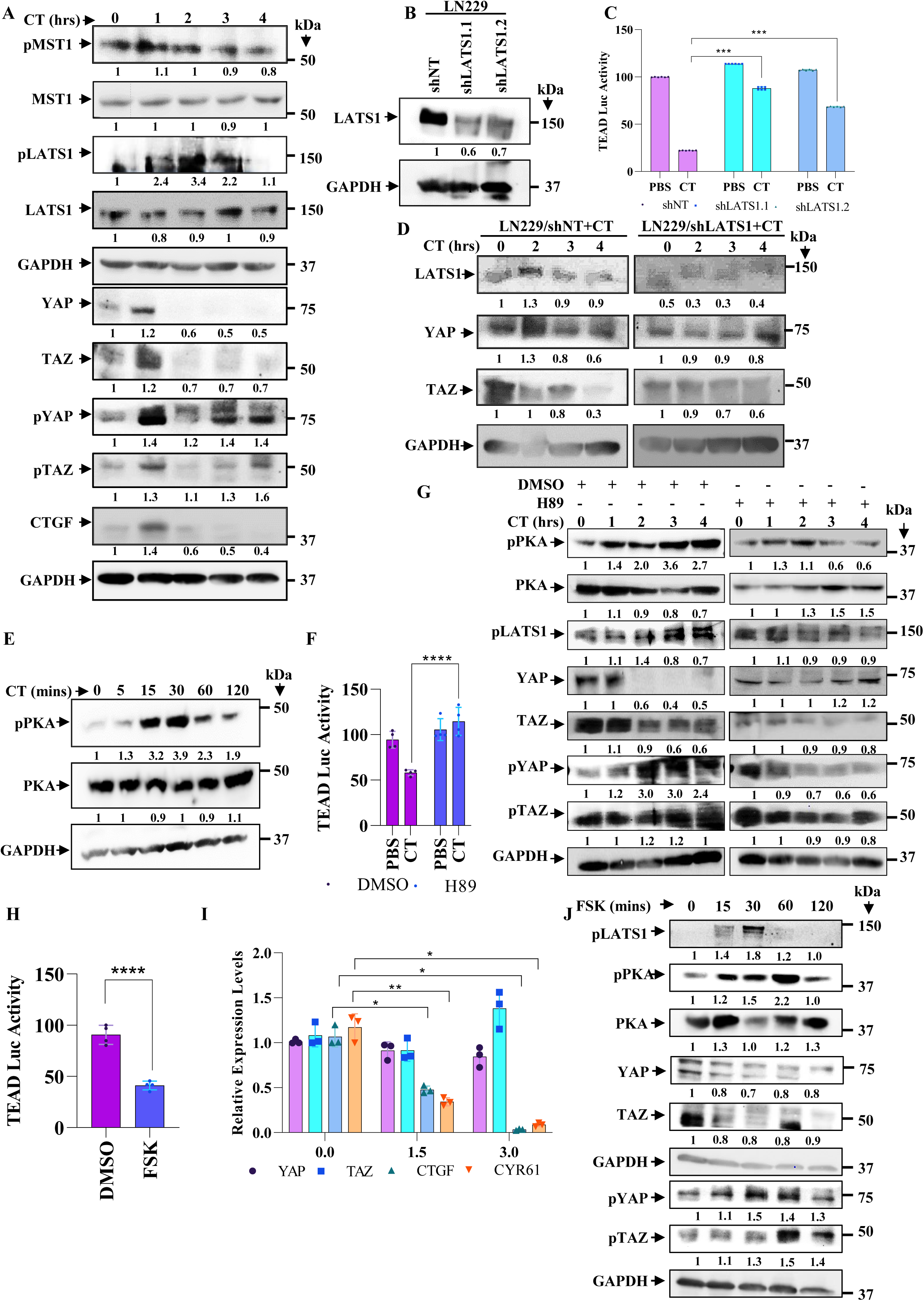
CT modulates YAP and TAZ in a cAMP-PKA-LATS1-dependent manner. **(A)** Western blot shows the effect of CT treatment on YAP, TAZ, and the upstream regulatory kinases MST1 and LATS1 in LN229 cells. (**B)** Western blot shows LATS1 knockdown using two shRNAs in LN229 cells. (**C)** LN229/shNT and LN229/shLATS1 cells were transfected with the TEAD luc construct, and the luciferase activity was determined after four hours of CT treatment. (**D)** Western blot shows the levels of YAP and TAZ upon CT treatment under shNT and shLATS1 conditions. **(E)** Western blotting shows the activation status of PKA in a time-dependent manner upon CT treatment. (**F)** LN229 cells expressing TEAD-luc were pre-treated with DMSO or H89 for 2 hours before CT treatment. The cell extracts were used to determine the relative TEAD Luc activity (two-way ANOVA; n=4/group). (**G)** The western blot shows the effect of CT on the PKA, LATS1, YAP, and TAZ in DMSO or H89-pretreated LN229 cells. (**H)** LN229 cells showing TEAD Luc activity in DMSO and FSK- (10 μM) treated cells (t test; n=4/group). (**I)** Expression of YAP, TAZ, CTGF, and CYR61 upon FSK treatment at the indicated time points (two-way ANOVA n=3/group). (**J)** Western blotting shows the levels of YAP and TAZ and their phosphorylated forms upon FSK treatment at the indicated time points.

The cAMP-dependent protein kinase A (PKA) axis has been reported to facilitate Hippo pathway activation through phosphorylation of LATS1 [**20, 21**]. Aptly, CT treatment of glioma cells resulted in a marked increase in phosphorylated PKA (pPKA) levels, without affecting the total abundance of PKA protein (**Figure 2E**). Importantly, pharmacological inhibition of PKA with H89 abrogated CT-mediated Hippo activation and suppression of cell proliferation (**Figure 2F, G; Supplementary Figure 2G, H**). As CT/CTR signaling is known to stimulate adenylate cyclase [**22**], we next examined intracellular cAMP levels and observed a significant elevation following CT treatment (**Supplementary Figure 2I**). Furthermore, the addition of forskolin (FSK), a direct activator of adenylate cyclase, recapitulated the effects of CT treatment on glioma cells (**Figure 2H, I, and J; Supplementary Figure 2J, K**). Collectively, these findings define a mechanistic framework in which CT/CTR signaling activates the Hippo pathway through a cAMP/PKA/LATS1 cascade, resulting in the suppression of YAP/TAZ transcriptional activity and inhibition of glioma cell proliferation.

### Glioma stem-like cells (GSCs) express elevated levels of CTR, and CT treatment inhibits their growth in vitro and GSC-initiated tumors in vivo

Glioma stem-like cells (GSCs), though representing a minor subpopulation within glioblastoma (GBM), are crucial for tumor initiation, maintenance, therapeutic resistance, and recurrence **[23]**. Patient-derived human GSCs, murine GSCs showed elevated expression of the calcitonin receptor (CTR) and YAP/TAZ compared to matched differentiated glioma cells (DGCs) (**Figure 3A**). Supporting their stem-like identity, GSCs from MGG8 and other models expressed higher levels of glioma reprogramming factors (SOX2, SALL2, and POU3F2) relative to their matched DGCs (**Supplementary Figure 3A**; [**24**]). Collectively, the high expression of CTR and YAP/TAZ underscores the vulnerability of GSCs to therapeutic strategies that activate the Hippo pathway. Indeed, CT treatment of human and murine GSCs activated the Hippo pathway in GSC lines, MGG4 (**Figure 3B, C, and D**), GB3 **(Supplementary Figure 3B, C, and D),** MGG8 (**Supplementary Figure 4A),** and DBT-Luc **(Supplementary Figure 4D).** Additionally, CT treatment inhibited the growth of human GSCs, MGG4 (**Figure 3 E, F, G, and H)**, GB3 **(Supplementary Figure 3 E, F, and G)**, MGG8 **(Supplementary Figure 4 B, C, and D)**, and murine GSC, DBT-Luc **(Supplementary Figure 4 E, and F)** as seen from the neurosphere growth assay and limiting dilution assay. We also demonstrate that GSCs are more susceptible to growth inhibition following CT treatment compared to a matched DGC (**Supplementary Figure 3H**). Further, MGG4 GSCs expressing the S127A-YAP mutant are found to be resistant to CT-mediated GSC growth inhibition due to inefficient TEAD-Luc inhibition (**Figure 3I)**, and persistent YAP abundance observed by confocal microscopy (**Figure 3J)** and continued proliferation (**Figure 3K)** in CT-treated cells. These results suggest that GSCs can be targeted specifically by CT-dependent activation of the Hippo pathway and inhibition of YAP/TAZ.

**Figure 3.**
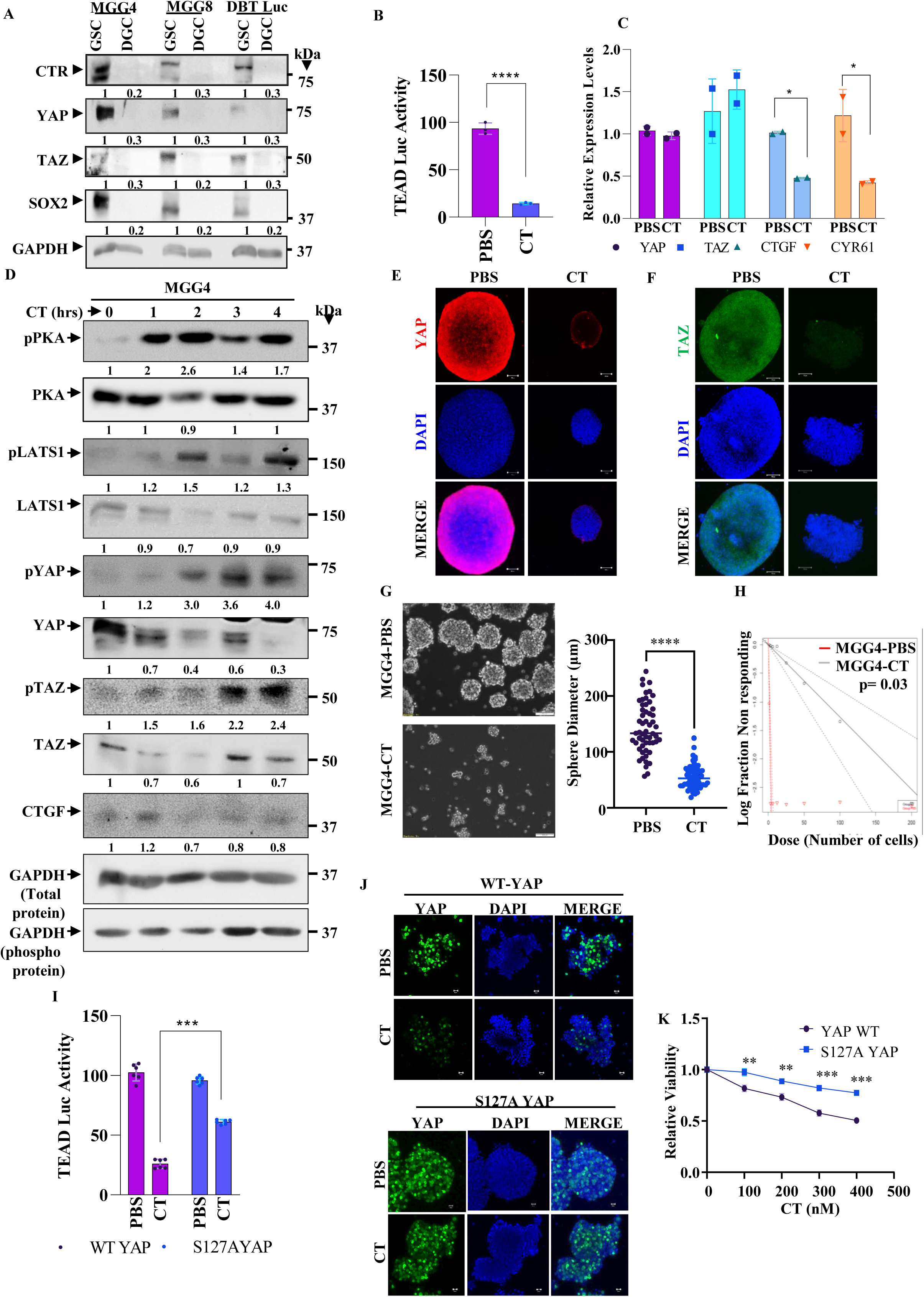
GSCs are better suited for CT-mediated growth inhibition. **(A)** Western blotting shows differential expression of YAP, TAZ, and CTR in GSC vs. DGC of MGG4, MGG8, and DBT-Luc. **(B)** TEAD-Luc activity in MGG4 cells following PBS or CT treatment (t test: n=3/group). **(C)** RT-qPCR analysis shows YAP, TAZ, CTGF, and CYR61 transcript levels upon CT treatment in the MGG4 cells (t test; n=2/group). **(D)** Western blot showing the effect of CT treatment on YAP, TAZ, and the upstream regulatory kinases MST1 and LATS1 in MGG4 cells. **(E, F)** Confocal microscopy analysis shows YAP and TAZ levels in MGG4 spheres with and without CT treatment. Red indicates YAP, green indicates TAZ, and blue indicates DAPI. Magnification x20, scale bar 100 µm. The merged images are shown for representation. **(G, H)** The sphere formation assay and the Limiting Dilution Assay show the effect of CT on the growth of MGG4 cells. **(I)** The plot shows TEAD Luc activity from LN229 cells expressing WT YAP or constitutively active YAP S127A upon treatment with PBS or CT (two-way ANOVA; n=6/group). **(J)** Confocal imaging shows the effect of CT on MGG4 spheres expressing WT-YAP or S127A YAP. **(K)** The graph shows the effect of CT on the viability of LN229 cells expressing WT or S127A YAP (two-way ANOVA; n=3/group).

Next, we investigated the ability of salmon CT to inhibit the growth of human and murine gliomas initiated by GSCs. Since salmon CT is a large peptide (32 amino acids) that cannot pass through the blood-brain barrier **[16],** we delivered salmon CT through the intranasal delivery route, an established route for the delivery of peptides and small molecules for the treatment of brain tumors and neurodegenerative diseases in mice **[17, 18]**. Intranasal delivery of CT inhibited the growth of glioma initiated by human (MGG4) and murine (DBT-Luc) GSCs and increased mouse survival (MGG4; **Figure 4 A, B, C, and D**)**, and (**DBT-Luc; **Supplementary Figure 5A, B, C, and D).** The sections derived from the MGG4-initiated small tumors formed in CT-delivered mice showed reduced staining for total YAP and TAZ (**Figure 4E and F; Supplementary Figure 5E and F**), increased pYAP/pTAZ (**Figure 4G**), increased pLATS1 with no change in total LATS1 (**Figure 4H and I**), and increased pPKA with no change in total PKA (**Figure 4J and K**). From these results, we conclude that CT inhibits GSC growth in vitro and glioma tumors initiated by GSCs through Hippo pathway activation in mouse models, thereby raising the possibility of using CT for intranasal delivery in GBM therapy.

**Figure 4.**
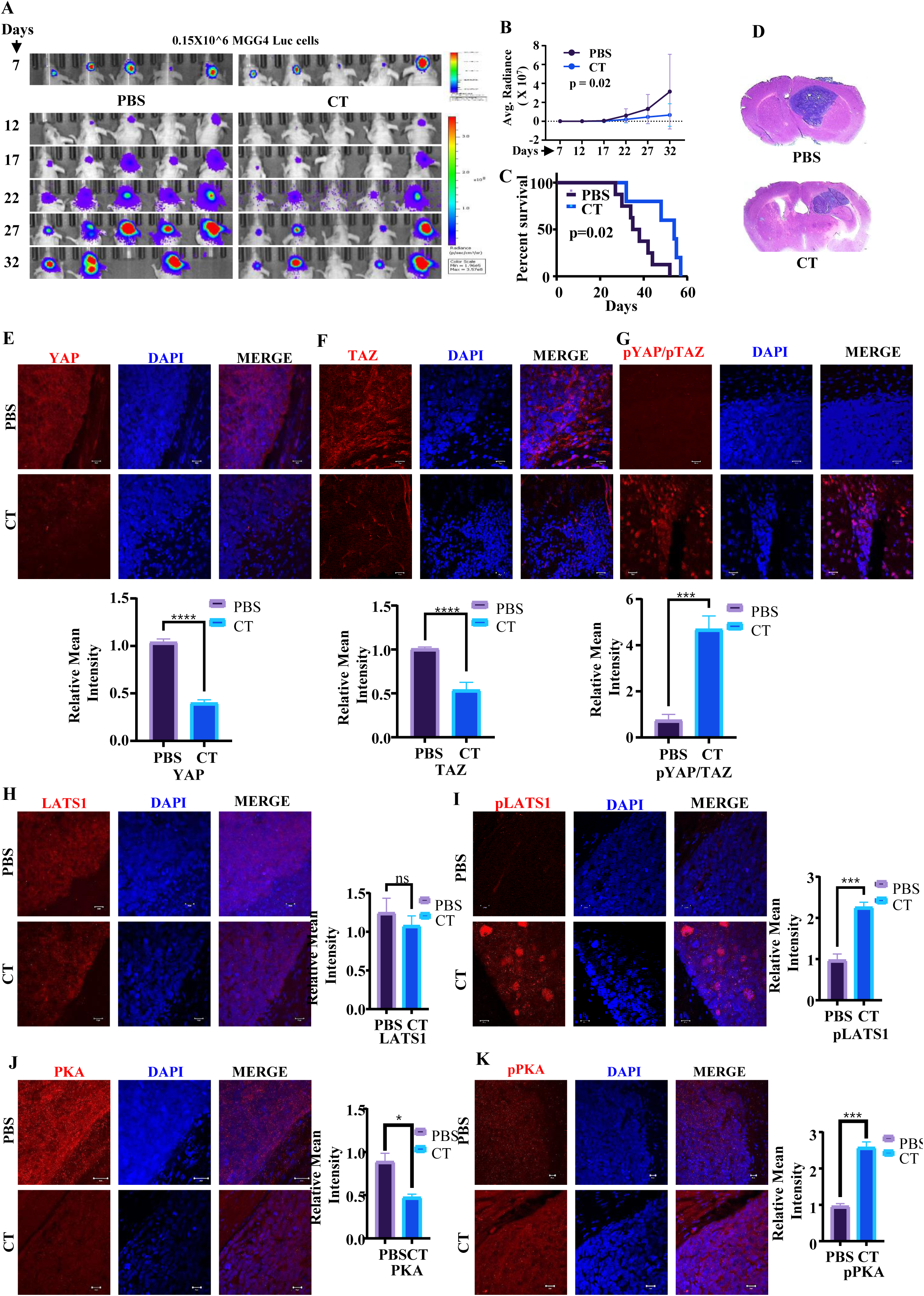
Intranasal CT administration inhibits MGG4-Luc GSC-derived tumors in nude mice. **(A)** 0.15×10^6^ MGG4-Luc GSCs were injected intracranially into the brain of nude mice. The tumor was allowed to grow for seven days, after which CT was administered intranasally at a dose of 2 IU/ animal. Representative images of the animals’ in vivo bioluminescence were taken every five days from the date of injection. (**B)** The average radiance efficiency between the PBS and CT-administered groups is plotted (two-way ANOVA; n=8 in PBS group and n=7 in CT-treated group). (**C)** The Kaplan-Meier graph shows the survival difference between the two groups of mice that were administered PBS or CT. (**D)** Haematoxylin and Eosin staining shows a larger tumor (depicted by dark blue colour) in animals administered with intranasal PBS compared to a smaller tumor in the CT-administered group. Confocal microscopy shows level of YAP **(E)**, TAZ **(F),** pYAP/TAZ **(G),** LATS1 **(H)** pLATS1 **(I)** PKA **(J)** and pPKA **(K)** levels in the PBS and CT-treated groups (Magnification-63x, scale bar-20µm).

We next examined the distribution of CT at the tumor site following intranasal delivery and the potential toxicities of CT in mice. Miacalcin (synthetic salmon calcitonin) is an FDA-approved drug for postmenopausal osteoporosis, as it inhibits osteoclasts **[19]**. C57BL/6 mice bearing GSC-initiated tumors were delivered with two different concentrations of Cy3-labeled CT through the intranasal route. Fluorescent imaging confirmed that Cy3-labeled CT reached the tumor (**Supplementary Figure 6A**). Additionally, fluorescence imaging of various dissected organs was limited to the brain only (**Supplementary Figure 6B).** Towards assessing the possible toxicity, mice were administered varying amounts of CT (2 IU, 4 IU, and 8 IU) intranasally daily for four weeks, and various parameters, including water consumption, food consumption, body weight, Heart rate, Ejection Fraction, Cardiac output, Covariance of heart rate (CV%), hematological and biochemical parameters, were measured at the end of each week (**Supplementary Figure 6C)** and found to have no significant difference in CT-treated mice. (**Supplementary Figure 6D, E, F, G, H, I, J, and K; Supplementary Table 3**). H&E staining of the major organs obtained from the CT-administered mice showed no histological alterations compared to those of control mice **(Supplementary Figure 7)**. From these results, we conclude that no adverse effects were observed in CT-treated mice.

### Molecular Dynamics (MD) simulations of patient-derived CTR mutants elucidate the mechanism behind their loss-of-function

Our results thus far show that WT CTR activates the Hippo pathway to inhibit glioma cell proliferation. Tumour-derived mutations in CTR result in loss of function and fails to inhibit the growth of glioma cells **[10]**. Out of the seven mutations, three (R45Q, A51T, and P100L) are on the extracellular domain (ECD), two (R404C and R420C) on the Helix-8 of intracellular domain (ICD), one (V250M) at the junction of transmembrane 3 (TM3) and Intracellular loop 2 (ICL2), and one (A307V) on TM5 (**Supplementary Figure 8A**). As expected, CTR with tumor-derived mutations failed to activate the Hippo pathway, as seen in their inability to inhibit luciferase activity from the TEAD-Luc construct (**Supplementary Figure 8B**).

To obtain conformation dynamics insights into the effects of the patient-derived mutations on the CTR functions, we performed all-atom MD simulations of the WT and mutant CTR systems. A representative 3D schematic of the MD system, with marked location of the seven mutants on the CTR, is shown **(Figure 5A)**. All systems under consideration are listed (**Supplementary Table 4**). With the WT system as a control, we compare the simulation data for different mutant systems and quantify the changes in 3D conformations and fluctuations to understand the impact of each mutation on the function of CTR.

**Figure 5.**
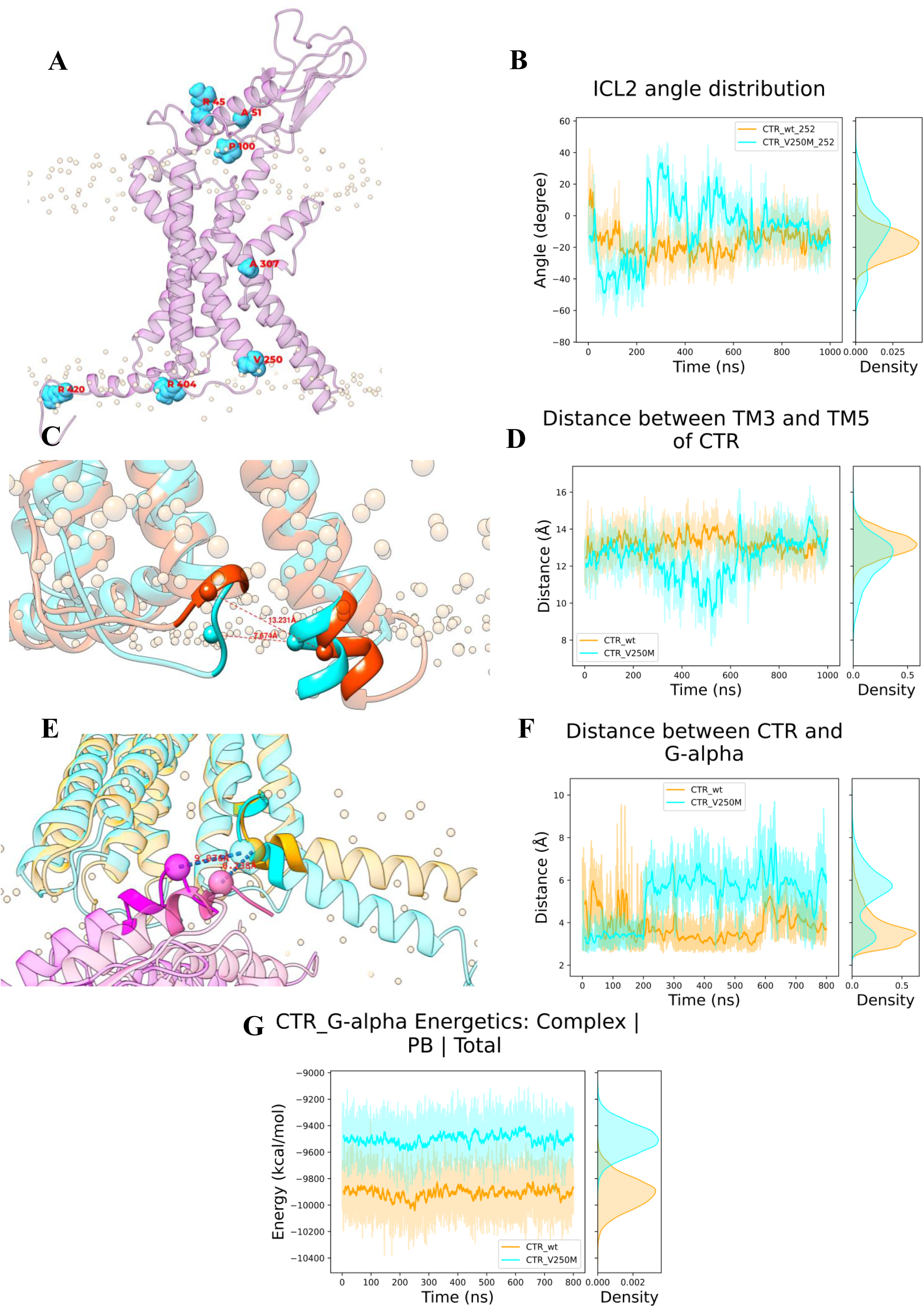
Illustration of mutant residue positions in CTR and differential conformation and interaction of V250M mutant compared to WT CTR. **(A)** Activated conformation of CTR with the position of seven patient-derived mutations is shown in cyan. **(B)** The plot shows ICL2 angle distribution in the WT CTR and V250M w.r.t. residue V252. **(C)** The figure shows the distance between the C-α carbon of the residue V/M250 in the transmembrane 3 (TM3) and residue K326 in the transmembrane-5 (TM5) in the WT CTR (orange) and V250M mutant (cyan). **(D)** The plot shows the distance between the Cα carbon of TM3 (V/M250) and TM5 (K326) between the WT CTR and the V250M mutant. **(E)** The plot shows the minimum distance between CTR (N396) and Gα (E392) in the WT CTR and V250M mutant. **(F)** The superimposed image shows the distance between the Cα carbon of residue N396 of CTR and residue E392 of Gα in the WT CTR (CTR in orange and Gα in pink) and V250M mutant (CTR in cyan and Gα in magenta). **(G)** Interaction energy between CTR and Gα in the WT CTR and V250M mutant. The red and blue colour lines denote the WT CTR and V250M mutant, respectively.

### V250M, an ICL2 mutant, affects ICL2 flexibility, TM5 conformation and Gα protein binding

Cryo-EM data (PDB ID: 6NIY) show that the 250th residue interacts with the G-protein **[12]**. Hence, we examined the impact of V250M on the flexibility of ICL2 in MD simulations of activated WT and the V250M systems. To quantify ICL2 flexibility, we defined two planes: a primary plane using C-α atoms of residues from TM3 (G241 and I248) and TM4 (R260), and a secondary plane involving C-α atoms of TM3 (I248), TM4 (R260), and an ICL2 residue as the fourth (V252) **(Supplementary Figure 8C).** Trajectory data revealed that the angle distribution between these planes is broader in the V250M mutant than in the WT CTR (**Figure 5B).** Since the V252 site is in the loop region, we wanted to check whether the flexibility is local to the V252 residue or the overall loop. Hence, we quantified the angle distribution, keeping other residues on the loop as the fourth site (F253, T254, or E255 of ICL2). This analysis also revealed a broader angle distribution for all the residues tested (**Supplementary Figure 8D, E, F**), thereby establishing that, overall, the ICL2 conformational dynamics has increased due to the V250M mutation. These changes in loop dynamics are functionally important as they can have a significant effect on the packing and conformation of proteins that could affect the protein-protein interaction network **[23–25]**. Indeed, investigation of the conformational difference between the activated states of the WT CTR and V250M mutant revealed that the distance between the C-α atom of residue at the 250th position in TM3 and the C-α atom of residue K326 in TM5 was reduced to approximately 10 Å (closest ∼7 Å) for the V250M mutant in contrast to ∼13 Å observed consistently in the trajectory of the WT CTR (**Figure 5C and D**). Furthermore, we observed that in the V250M mutant, TM5 bends towards

TM3 in the intracellular region, which is not observed in the WT CTR (**Figure 5C**). This conformational alteration is particularly significant because the intracellular segments of TM3 and TM5 are directly involved in interactions with Gα protein, suggesting that the V250M mutation could potentially impact the interaction between CTR and Gα protein.

We also investigated the impact of the observed conformational changes between WT CTR and V250M mutant on the interaction between Gα and CTR by measuring the distance between C-α atoms of specific residues CTR (N396) and Gα (E392). We find this distance to be smaller in the WT CTR than in the V250M mutant, and the Gα is seen slightly shifted out of the binding site for the V250M mutant (**Figure 5E, F**). This suggests that the V250M mutant may have a weaker binding affinity for Gα than the WT CTR. Furthermore, our examination using MMPBSA-based interaction energy calculations **[26, 27]** showed that the interaction energy between CTR and Gα is lower in the WT CTR than in the V250M mutant (**Figure 5G**). This further corroborates that V250M mutation weakens the interaction between CTR and Gα and consequently could affect the downstream signaling.

### C terminal Helix-8 mutants, R404C and R420C, show reduced membrane Interactions

The C-terminal mutation **R404C** is located within the central segment of Helix-8, while R420C resides within the unstructured C-terminal region following Helix-8 (**Figure 5A and Supplementary Figure 8A)**. To analyze the impact of these mutations, we calculated the angle between Helix-8 (axis joining C-α atoms of N396 and N414) and membrane normal (Z-axis). We find that the Helix-8 moves away from the membrane plane for the R404C mutant (**Figure 6A and B**) as well as for the R420C mutant **(Supplementary Figure 9A and B)** as compared to WT CTR. We also measured the residue 404 occupancy with PIP2 lipids on the membrane and found that R404 of WT CTR interacts closely with the PIP2 lipid with an occupancy of 94.68% and an average proximity of 2.5-3.0 Å. On the other hand, the distance is increased to 6-8 Å in the R404C mutant with a significantly lower occupancy of 5.11% (**Figure 6C**). Similarly, the distance between residue 420 and the PIP2 lipid is approximately 4-6 Å in the R420C mutant with an occupancy of 62.16% compared to around 2.5-3 Å in the WT CTR with a 99.98% occupancy (**Supplementary Figure 9C**). We also show the molecular-scale interaction profile for the PIP2-Helix 8 complex from the bottom view and side view for the R404C mutant (**Supplementary Figure 9D and 9E, respectively**) and for the R420C (**Supplementary Figure 9F and 9G, respectively**). While the front view (**Supplementary Figure 9A)** revealed that the Helix-8 had swivelled away from the membrane with significantly reduced membrane contact, the bottom and side views clearly show the loss of interaction with the PIP lipids. As seen for the R404C mutant (**Supplementary Figure 9D and E, respectively**) and for R420C (**Supplementary Figure 9F and G, respectively**), in the wild type, the Arginine interacts with the PIP2 lipid, while the mutant Cysteine moves away from PIP2 lipids with its side chain facing outwards. It has been demonstrated both experimentally [28] and through simulation studies [29] that the interaction between anionic lipids, such as PIP2, and the basic residues of GPCRs promotes/stabilizes the active conformation of GPCRs, thereby strengthening the interaction between basic residues at the interface of GPCRs and Gα. An equivalent interaction is observed between Arginine residues at the 404^th^ and 420^th^ positions and PIP2 lipids, and this interaction is lost when the Arginines are mutated to Cysteines. Hence, the Arginine to Cysteine mutation at 404 or 420 locations is likely to weaken the CTR interaction with G-protein complexes, possibly affecting CTR activity.

**Figure 6.**
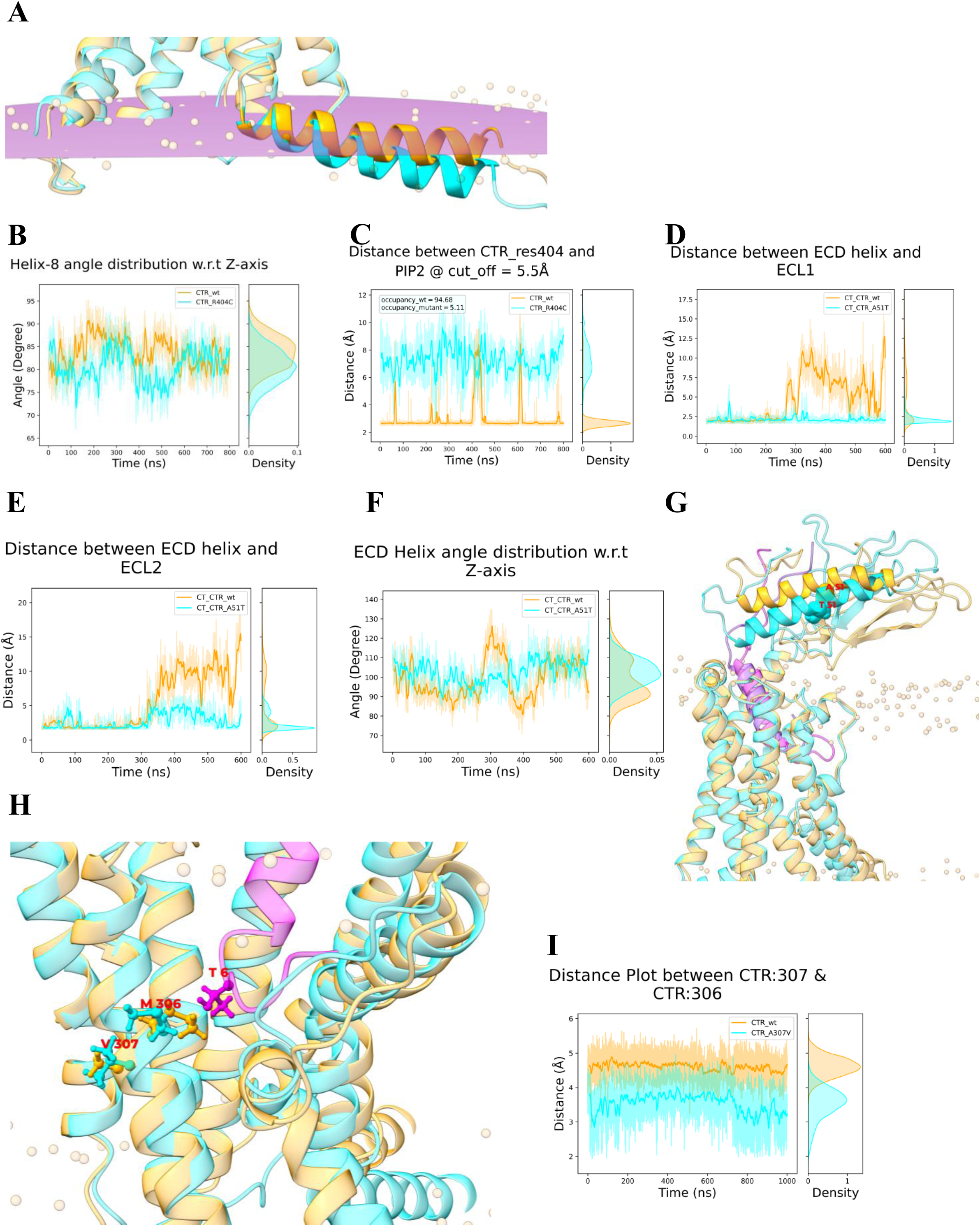
Conformational shift and differential interaction due to mutant CTRs R404C, A51T, and A307V with membrane lipid, ECLs, and CT, respectively, compared to WT CTR. **(A)** The superimposed image shows the orientation of the 8^th^ helix in the WT CTR (orange) and R404C mutant (cyan) CTR. **(B)** The plot shows the difference in the angle distribution of the 8^th^ helix with the z-axis in the WT CTR and R404C mutant. **(C)** The plot shows the interaction of the 404_th_ residue with PIP2 lipids of the membrane in the WT CTR and R404C mutant. **(D)** The superimposed image of WT CTR (orange) and A307V mutant CTR (cyan) shows the differential orientation between residues A/V307:CTR, M306:CTR, and T6:CT between WT CTR and mutant systems. **(E)** The plot shows the minimum distance between the side chain of residue A/V307 and M306 for WT CTR and the A307V mutant. **(F)** The plot shows the minimum distance between the ECD helix and ECL1 loop for WT CTR and A51T mutant when bound to CT. **(G)** This plot shows the minimum distance between the ECD helix and ECL1 loop for the previously mentioned systems. **(H)** The plot shows the ECD helix angle with respect to the Z-axis for the systems, the same as Figure F. **(I)** The superimposed image shows the ECD orientation between WT CTR (orange) and A51T mutant (cyan) when bound to CT (pink).

### ECD mutants, R45Q, A51T, and P100L, affects ECD and ECL1/ECL2 interaction

Next, we investigated the three ECD mutations, R45Q, A51T, and P100L (**Figure 5A and Supplementary Figure 8A)**. All three mutants induce noticeable structural changes in CTR-ECD that could have functional implications. The ECD with a long alpha-helix (F39-Q60) assumes a lid-like feature in the CTR. Since ECD conformations dictate CT recruitment and binding, we decided to focus on how the ECD alpha-helix interfaces with the ECL1 (P207 to P216) or ECL2 (F285 to H296) of the TM region of the protein for the WT CTR and for the ECD mutants. To achieve this, we calculated the minimum distance between the ECD-helix and the ECLs. Also, much like the Helix-8, we aimed to characterize the helix orientation relative to the membrane normal. Analyses show that for the A51T mutant, the ECD closes in on the TM region of CTR (around 2.0 - 4.0 Å) and acts like a tight lid, whereas the ECD has an expansive breathing motion in the WT CTR (2.0 - 20.0 Å). **Figures 6D and 6E** show the minimum distance distribution between ECD-ECL1 and ECD-ECL2, respectively, for both WT CTR and A51T mutant. Concomitantly, the WT CTR is also observed to have higher fluctuations in the helix angle than the A51T mutant (**Figure 6F).** Similar behaviour, though to a lesser extent, is observed for R45Q (**Supplementary Figure 10A, B, and C)** and P100L **(Supplementary Figure 10D, E, and F).**

Distance and angle analysis of the ECD helix, as well as conformational dynamics insights from the trajectories (**Figure 6G for A51T, Supplementary Figure 10G and H for R45Q and P100L, respectively)** clearly reveal that the ECD is more dynamic and flexible and favours an open conformation for WT CTR in the ligand-bound state compared to the mutants. Another interesting insight that we obtained from our simulation studies is that the ECD exhibits a closed-lid state when WT CTR is not bound to the CT ligand (data not shown). It appears that the binding of CT induces spatial rearrangements in the protein, disrupting the ECD-helix interactions with ECL1 and ECL2, which leads to a highly breathable and open ECD. ECD mutants, on the other hand, don’t disrupt these cross-interactions. Even though we observe conformational rigidity of ECD-ECLs in mutants structurally, the origin of this change could be a result of a combination of multiple short- and long-range weak interactions, as it is not straightforward to directly single out a specific set of interactions that induce this change.

### TM5 mutant, A307V, weakens the interaction between CT and CTR

CT binding to WT CTR induces conformational changes in CTR, with possible implications in downstream signaling **[7]**. Next, we studied the impact of ECD-juxtaposed residue in TM5 (A307V) on CT binding to WT CTR. We found a subtle effect of mutation on the strength of CTR-CT binding. The sixth residue of CT (T6) interacts with residue M306 of WT CTR, and there is a weaker interaction between the side chains of A307 and M306 on TM5 (**Figure 6H**). When the A307V mutation occurs, due to the increased reach of Valine, the interaction strength between V307 and M306 increases as the minimum distance between their side chain decreases (**Figure 6I**). It is likely that this stronger interaction between V307 and M306 weakens the interaction between M306:CTR and T6:CT. Such hallmark subtle interactions are often widespread in the GPCR systems [**30**], and high-resolution AAMD simulations usually help unravel such networks of interactions, as shown above.

Together, molecular simulation data at atomic resolutions helped us understand the effect of mutants on the CTR signaling pathway from both structural and conformational dynamics perspectives. For example, our data show that a mutation in the intracellular region of the protein (V250M) significantly enhances the ICL2 flexibility, which in turn can affect G-protein binding. We also found that R404C and R420C mutations in the intracellular region remarkably reduce the interaction strength between helix-8 and the membrane, which could potentially impact the active CTR population and its interaction with G-proteins. Our analyses of mutants located in the ECD (R45Q, A51T, and P100L) indicate that these mutants shift the ECD towards an alternate configuration, potentially influencing CT retention and binding. The A307V mutant on TM5 exhibits an enhanced interaction with CT-binding residues, which may, in turn, weaken the CT-CTR signaling. In conjunction with the available experimental data, our in silico conformational landscape investigations of the WT and mutant CTR systems (in the presence of CT and G-proteins) can be used to establish a hypothesis towards molecular mechanisms of CTR activation in health and disease states.

## Discussion

In our previous study, the neuroactive ligand-receptor interaction pathway emerged as one of the prominently mutated pathways with prognostic value for GBM patient survival. Notably, CTR was identified as the most frequently mutated gene, with mutations detected in 3.0% of GBM patients; such patients with mutated CTR had poor survival outcomes. Interestingly, CT treatment of glioma cells with WT CTR inhibited the growth of glioma in vitro and in vivo **[10]**. However, the underlying mechanism of this tumor suppressor function of the CT/CTR axis has remained elusive. In this study, we found that the CT/CTR axis promotes the degradation of YAP/TAZ, well-known oncogenic transcription factors, by activating the Hippo pathway. Hippo pathway activation required cAMP/PKA/LATS1 signaling pathway. Intranasal delivery of salmon CT inhibited glioma growth efficiently. Dynamis simulation studies revealed defects in the interactions between key residues in mutant CACLR upon CT binding, unlike WT CTR.

The Hippo pathway, found to be conserved among higher vertebrates, serves as a critical signaling pathway involved in regulating organ size and maintaining tissue homeostasis **[2]**. Dysregulation of the Hippo pathway is reported in multiple cancers, including GBM **[31, 32]**. Our investigation revealed that treatment of GBM cells with CT led to the inhibition of YAP and TAZ proteins through LATS1-dependent phosphorylation. We also found that the cAMP-PKA-LATS1 pathway mediated the regulation of YAP and TAZ by the CT/CTR axis. This axis of regulation aligns with findings in muscle stem cells, where the CTR-PKA-LATS axis similarly regulates YAP and TAZ activity, thereby contributing to the maintenance of muscle stem cells.

It is now well established that GSCs, which form small proportions of the tumors, are the tumor-initiating cells that drive tumor growth and impart resistance to conventional therapy **[33]** underscoring the need to identify targets within the therapy-resistant pool of GSCs to develop add-on therapies that increase the efficacy of the current treatments. Interestingly, our investigation revealed elevated expression levels of CTR in patient-derived GSCs compared to matched differentiated glioma cells (DGCs). We also found high levels of YAP and TAZ in patient-derived GSCs. Involvement of YAP and TAZ in preserving the stemness and plasticity of GSCs in GBM and other cancers **[34, 35]**. Thus, the high levels of CTR and YAP/TAZ observed in GSCs make them suitable candidates for CT treatment in glioma therapy. Our results show that treatment of GSCs with CT resulted in a pronounced inhibition of GSC growth. Furthermore, we found, based on whole-tumor transcriptome studies, that CTR transcript levels are significantly higher in the mesenchymal gene expression subtype, but there is no difference between G-CIMP, IDH mutation, and MGMT promoter methylation status (**Supplementary Figure 11**). This suggests that mesenchymal GBM may be more suitable for CT-based therapy. Further, intranasal administration of CT demonstrated an efficient reduction in the tumor volume initiated by both patient-derived human GSCs and murine GSCs in an intracranial orthotopic mouse glioma model. Collectively, these findings underscore the promising therapeutic potential of CT in targeting tumor-initiating cell populations.

Structural and dynamic insights from microseconds-long simulation studies provide information about the possible activation mechanisms of CTR and offer a rationale for the loss of function in patient-derived mutants. Class B GPCRs, such as CTR, have a characteristically large extracellular domain that has evolved to detect and bind the ligands in a tightly regulated manner. Recent alanine scanning mutagenesis data on ECL1[**14**] and ECL2/ECL3 [**13**] from Sexton and co-workers clearly exhibit how the ECL residues control the binding and signaling of distinct CT. Interestingly, our data show that patient-derived mutant systems (R45Q, A51T, and P100L) exhibit a remarkably different orientational conformation of the ECD, with compromised ECL-ECD interactions that affect CT encapsulation. From this, we may infer that CTR-ECD, through its interaction with the ECLs, seems to act like a well-designed lid for CT recognition. Also, our data from the A307V mutation studies clearly reveal that CT-binding triggers a subtle packing rearrangement in the membrane-embedded TM region, which transfers the information to the G-protein binding ICL region of the CTR [**36–38**]. Several reports exist in GPCR literature on the crucial role of ECLs [**13, 14**] and ICLs [**39**]. V250M, a mutation on the ICL2 in CTR, causes profound changes in the ICL2 flexibility that affect the binding interface with the alpha-subunit of the G-proteins. Besides the altered interaction with CT and G-proteins, our simulation provides a structural origin of the loss of function by indicating a reduced interaction with the anionic lipids of the membrane, and the CTR with the mutations, i.e., R404C and R420C, on Helix-8 and further down, respectively – a factor that could compromise the CTR function. Beyond structurally connecting the CT/CTR signaling axis and providing mechanistic insights into the signaling cascade, the conformational landscape presented by the simulation studies could also be used to rationally design more effective CT-mimics as therapeutic agents to restore the YAP-TAZ activity.

We propose a model (**Figure 7**) that describes the Hippo pathway activation by the CT/CTR axis and the mechanism behind loss-of-functions by patient-derived CTR mutants. Overall, this study provides significant insights into the mechanism underlying the growth inhibitory pathway of the CT/CTR axis in GBM, primarily through activating the Hippo signaling pathway. Furthermore, we have substantiated the therapeutic significance of intranasal CT treatment in glioma. The salmon calcitonin utilized in this research is an approved drug for women with post-menopausal osteoporosis, intended to maintain bone strength. Repurposing salmon calcitonin for GBM patients with wild-type CTR holds promise for inhibiting glioma growth.

**Figure 7.**
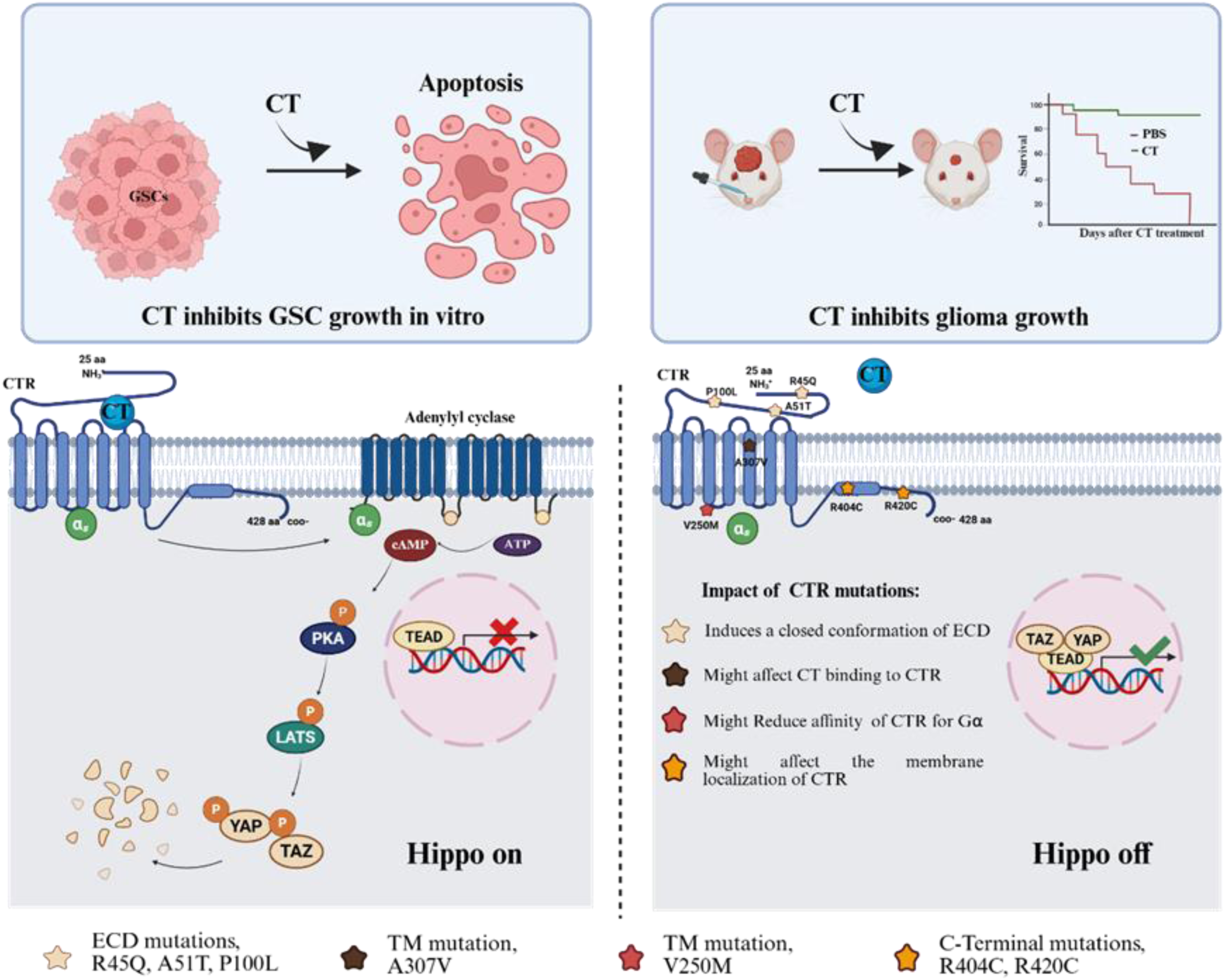
The proposed model describes the Hippo pathway activation by the CT/CTR axis, repurposed use of salmon CT for GBM therapy, and the mechanism behind loss-of-functions by patient-derived CALCR mutants.

## Author contributions

**JG, PU, KRP, AC, LG, SS, JC, AS, KS**

JG planned and carried out all the experiments; PU carried out all the structural analyses; KRP and JG carried out animal experiments; AC carried out the RPPA analysis; LG helped with the Confocal microscopy experiments, toxicity studies, animal experiments, and bioavailability experiments; SS and JC helped with the Cy3 labeling of CT. AS and KS prepared the manuscript, planned the study, and led the project.

## Data Availability statement

Any high-throughput data generated will be submitted to online depositories, and the same will be intimated to the journal office

## Acknowledgments

We acknowledge the shRNA consortium (Dr. Subba Rao), IISc, India, for shRNA constructs. The results published here are, in whole or in part, based upon data generated by The Cancer Genome Atlas (TCGA) pilot project established by the National Cancer Institute (NCI) and the National Human Genome Research Institute (NHGRI). Information about TCGA and the investigators and institutions that constitute the TCGA research network can be found at http://cancergenome.nih.gov. JG acknowledges the IISc fellowship, PU thanks CSIR for the graduate fellowship (**09/079(2868)/2021-EMR-I**), and KRP acknowledges DBT for the fellowship. KS acknowledges CEFIPRA, DBT, DST, and CSIR (Govt. of India) for research grants. Infrastructure supported by DST FIST, DBT-IISc partnership program, and UGC is acknowledged. AS acknowledges the financial support from the Indian Institute of Science-Bangalore and the high-performance computing facility “Beagle” that was set up from grants by a partnership between the Department of Biotechnology of India and the Indian Institute of Science (IISc-DBT partnership programme). AS thanks the DST for the National Supercomputing Mission grants (DST/NSM/R\&D-HPC-Applications/2021/03.10, DST/NSM/R\&D-HPC-Applications/ Extension Grant/ 2023/27). AS also thanks the DST for the FIST program sponsored by the Department of Science and Technology. AS would also like to thank the Teams Science Grant from the DBT-Wellcome Trust India Alliance (Grant number: IA/TSG/21/1/600245). AS also thanks the DBT National Network Project (NNP) grant (BT/PR40323/BTIS/137/78/2023) and the Matrics grants (MTR/2023/001040) from the Science and Engineering Board (SERB), India.

## Abbreviations

CT: Calcitonin
CTR: Calcitonin Receptor
YAP: Yes-Associated Protein
TAZ: Transcriptional coactivator with PDZ-binding motif
LATS1: Large Tumor Suppressor Kinase 1
PKA: Protein Kinase A
cAMP: cyclic Adenosine Monophosphate
GSC: Glioma Stem-like cells
DGC: Differentiated Glioma cells

## Supplementary file

### Other Methods and Materials

#### Cell lines used

We received the human tumor-derived GSCs MGG4 and MGG8 as generous gifts from Dr. Wakimoto at Massachusetts General Hospital, Boston, USA. The neurosphere, GB3, was established in our lab using GBM tumor samples. The DBT-Luc cells were provided by Dr. Dinesh Thotala at Washington University in St. Louis, St Louis, MO. The LN229 cell line was procured from ATCC (# CRL-2611, https://www.atcc.org/products/crl-2611), and the T98G cell line from ATCC (# CRL-1690, https://www.atcc.org/products/crl-1690). The U87 cell line was purchased from the ECACC (#89081402). All the cell lines were procured from their respective sources within the last year, with periodic authentication procedures conducted as necessary. Mycoplasma contamination was routinely monitored and confirmed absent using the Ezdetect PCR Kit for Mycoplasma Detection (HiMedia).

#### Reverse phase protein array analysis (RPPA) for the identification of differential protein expression upon CT treatment

CT-treated and untreated LN229 cells were lysed using RPPA lysis buffer. The resulting lysates were serially diluted in five twofold steps using the same lysis buffer and then printed onto nitrocellulose-coated slides using an Aushon Biosystem 2470 arrayer. Afterwards, the slides were probed with 441 validated primary antibodies, followed by detection using suitable biotinylated secondary antibodies. The slides were scanned, analyzed, and quantified using Array-pro Analyzer software (Media Cybernetics) to obtain spot intensity data (level 1 data). Signal visualization was achieved through a secondary streptavidin-conjugated HRP antibody and DAB colorimetric reaction. The list of the 441 antibodies utilized is provided at https://www.mdanderson.org/research/research-resources/core-facilities/functional-proteomics-rppa-core/education-and-references.html.

#### Neurosphere culturing

GSCs were isolated from GBM tumor tissue following a series of steps. Initially, the tissue was dissected and treated with trypsin, followed by the addition of a trypsin inhibitor. The treated tissue was mechanically dissociated and filtered to remove any debris. The resulting filtrate was cultured in ultra-low attachment plates in a specialized stem cell medium suitable for neurosphere formation. These neurospheres were cultured in a Neurobasal medium supplemented with various growth factors and supplements, including l-glutamine, heparin, B27 supplement, N2 supplement, rhEGF, rhFGF-basic, and antibiotics. After seven days of culture, the neurospheres were chemically dissociated into single-cell suspensions using a NeuroCult Chemical Dissociation Kit for subsequent replating.

#### Differentiation of GSCs to DGCs

To induce the differentiation of GSCs into DGCs, mature neurospheres were harvested from ultra-low attachment plates (Corning, USA). These neurospheres were transferred onto standard cell culture dishes and cultured in Dulbecco’s Modified Eagle Medium (DMEM) supplemented with 10% fetal bovine serum (FBS) and antibiotics, as previously described, for 15–20 days **[1]**.

#### Reprogramming of DGCs

After cultivation in DMEM supplemented with 10% FBS, the differentiated GSCs and GBM cell lines were subjected to two rounds of centrifugation in phosphate-buffered saline (PBS) to eliminate residual FBS. Subsequently, they were seeded onto ultra-low attachment plates and cultured in Neurobasal medium supplemented with the previously mentioned supplements and antibiotics for 7–10 days.

#### Limiting Dilution Assay

Sphere formation assays were conducted by plating 1, 10, 50, 100, and 200 individual glioblastoma stem cells (GSCs) per well in 10 wells each of a 96-well plate. Over the subsequent 5–7 days, the presence or absence of sphere formation was monitored. The count of wells where sphere formation did not occur was recorded and plotted against the initial number of cells per well. Utilizing the Extreme Limiting Dilution Assay (ELDA) software, which is accessible online, and the data analysis was performed (https://bioinf.wehi.edu.au/software/elda/).

#### Lentivirus preparation and transduction of cells

HEK-293T cells were seeded onto poly-l-lysine-coated cell culture dishes of either 60 mm or 90 mm diameter. Upon reaching 60–70% confluency, the cells were transfected with shRNA plasmid alongside helper plasmids psPAX2 and pMD2.G using Opti-MEM medium (Invitrogen #22600-050) and Lipofectamine 2000 (https://www.thermofisher.com/order/catalog/ product/12566014; Invitrogen) facilitated the transfection process. Following a 6-hour incubation period post-transfection, the Opti-MEM medium was replaced with fresh DMEM supplemented with 10% FBS. After 60 hours, the supernatant containing the transfected cells’ contents was harvested into 15 ml Falcon tubes, subjected to centrifugation at 5000 rpm for 10 minutes, filtered through a 0.45 μm filter, and finally stored at –80°C for future applications. The shRNA clones used in the study were taken from the Sigma human TRC shRNA library. For the knockdown of CALCR, the shRNA clones used were TRCN0000008162 and TRCN0000008163. The shRNA clones TRCN0000196268 and TRCN0000195739 were used for the knockdown of LATS1.

#### Luciferase and beta-galactosidase (β-gal) assay

Cells were seeded into six-well plates and co-transfected with a luciferase reporter construct and pCMV-beta gal (as a transfection control) with the help of Lipofectamine 2000 reagent following the manufacturer’s instructions. After 24 hours of transfection, cells were treated with CT for 4 hours before being harvested. Cells were lysed using the reporter lysis buffer (Promega), and the protein concentrations were estimated using the Bradford assay reagent (Bio-Rad). Subsequently, 10 µg of protein was added to 50 µl of luciferase assay reagent (LAR) to evaluate luciferase activity, which was later normalized to β-galactosidase activity units.

#### RNA isolation, cDNA conversion, and Real-Time quantitative qPCR

Total RNA from cells was isolated using TRI reagent following the manufacturer’s protocol. Subsequently, the RNA quality was determined by electrophoresis on a 2% MOPS-formaldehyde gel, and RNA was quantified using NanoDrop. 2 µg of total RNA was used for cDNA conversion using a high-capacity cDNA reverse transcription kit. The cDNA was diluted to a final concentration of 10 ng/µl. Real-time quantitative PCR was carried out using ABI Quant Studio 5 and 6 instruments, using the DyNAmo Flash SYBR Green qPCR kit. Gene-specific primer sets were used under defined cycling conditions for the reaction. Housekeeping genes such as GAPDH, ACTB, 18S rRNA, RPL35a, and ATP5G1 were used as human gene expression analysis references. Gene expression was calculated using the ΔΔCT method, transformed to a log2 ratio, and converted to the absolute scale for visualization. The primers used for RT-qPCR are shown below.

#### List of primers used in the study

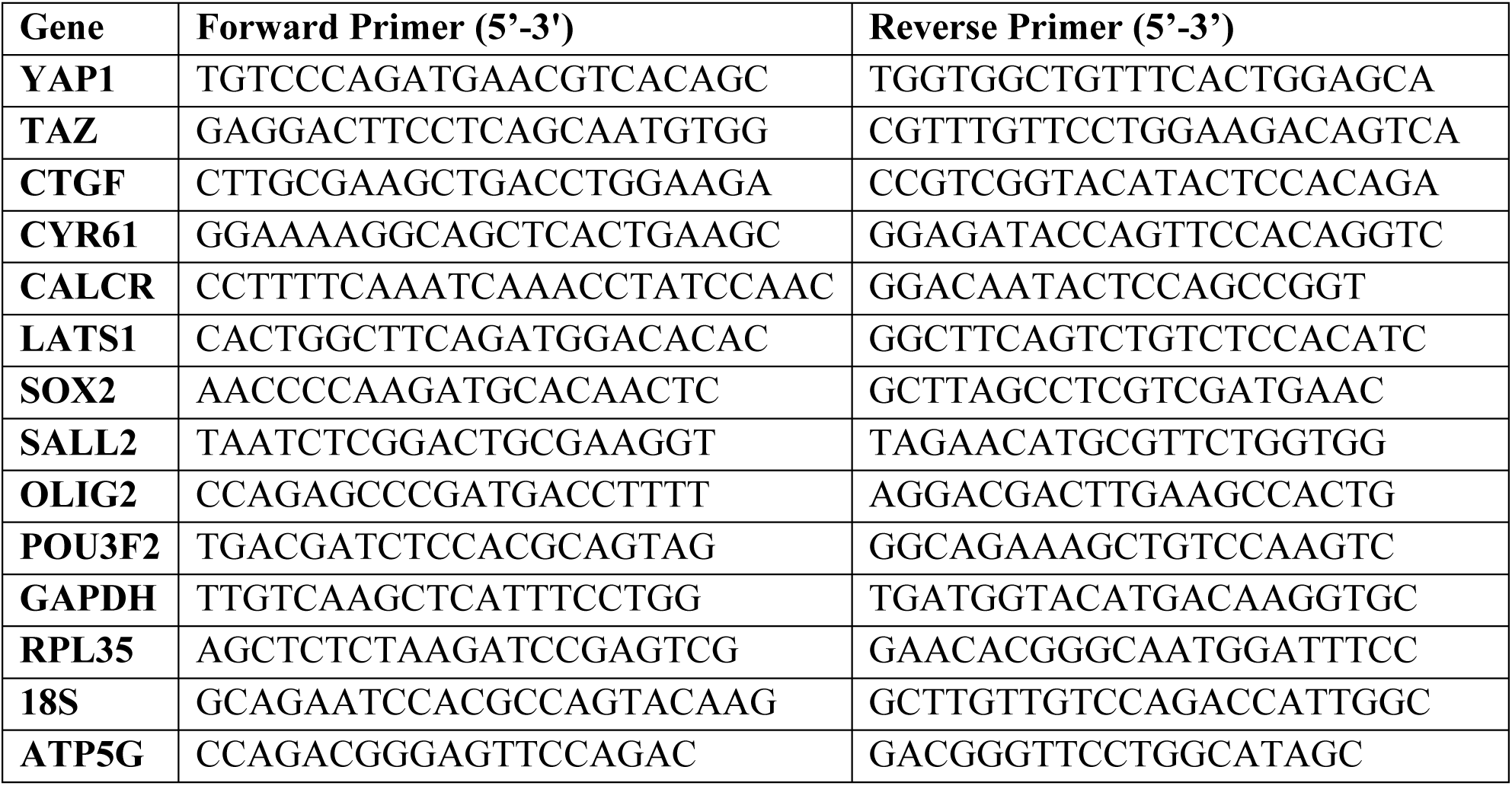

**Western Blotting**

For Western blot, cell pellets were lysed in RIPA lysis buffer containing specific protease inhibitors. Following lysis, the cells were centrifuged, and the protein supernatant was collected. The protein concentration in the sample was estimated using Bradford’s reagent, with a BSA curve serving as a standard reference. Equal protein amounts were added with loading dye, denatured at 90°C for ten minutes, and loaded onto an SDS PAGE gel. The gel electrophoresis was conducted for approximately 8 hours, with resolving and stacking gels prepared at appropriate concentrations. Proteins were then transferred onto PVDF membranes using a semi-dry transfer apparatus. After blocking with skimmed milk, the membrane was probed with primary antibodies overnight at 4°C, followed by washing three times and incubation with secondary antibodies. Subsequently, the blot was visualized using chemiluminescent reagents and imaged using a BIO-RAD chemidoc imaging system.

#### Antibodies and reagents used

The primary antibodies used in the study were purchased from the following companies: CTR (Invitrogen PA5-25594, 1:1000), YAP (CST 14074S, 1:1000), TAZ (CST 4883S, 1:1000), pYAP (CST 4911S, 1:1000), CTGF (CST 86641S, 1:1000), LATS1 (CST 9153S, 1:500), pLATS1 (CST 8654S 1:500), MST1 (CST 3682S, 1:1000), pMST1 (CST 49332S), GFP (CST 2956S), PKA (CST4782S, 1:1000), pPKA (CST 5661S, 1:1000), SOX2 (Abcam ab92494, 1:2000), GAPDH (SIGMA G8795, 1:25000). Goat anti-rabbit IgG (H+L) secondary HRP conjugate (Invitrogen# 31460, WB 1:5000), Goat anti-mouse IgG (H+L) secondary HRP conjugate (Invitrogen# 31430, WB 1:5000), goat anti-rabbit IgG (H+L) highly cross-adsorbed secondary antibody, Alexa Fluor 488 (Invitrogen, Cat# A-11034), goat anti-rabbit IgG (H+L) highly cross-adsorbed secondary antibody, Alexa Fluor 594 (Invitrogen, Cat# A-11037). All the Alexa Fluor antibodies were diluted at 1:500 for ICC and IHC. Salmon calcitonin (Unicalcin 100 IU) was purchased from a chemist. The list of small-molecule inhibitors used in this study is given below.

#### List of inhibitors used in the study

| **S.No.** | **Inhibitor name** | **Target molecule** | **Catalog Number** |
| --- | --- | --- | --- |
| **1.** | TRULI | LATS | HY138489 (Medchem Express) |
| **2.** | H89 | PKA | B1427 (Sigma Aldrich) |
| **3.** | Forskolin | Adenylyl cyclase | F6886 (Sigma Aldrich) |

**Apoptosis**

The Annexin V Apoptosis Detection Kit from BD Biosciences was used to quantify apoptotic cells. Followed by CT treatment for 72 hours, cells were washed twice with cold PBS and then suspended in a binding buffer at a concentration of 1 × 10^6 cells per ml. Subsequently, 100 μl of this suspension (equivalent to 1 × 10^5 cells) was transferred to a 5-ml culture tube, and 5 μl of annexin V was added. Following a 15-minute incubation at room temperature in the dark, an additional 400 μl of binding buffer was added to each tube. The cells were then analyzed using a flow cytometer (FACS Verse, BD Biosciences) within 1 hour of staining.

#### Colony Formation

LN229 cells were seeded at a density of 1×10^3 cells per well in a 6-well plate. After 24 hours, the cells were subjected to treatments with either PBS or CT, which were subsequently replenished every other day for 10 days. On the 11th day, the cells were fixed, stained using crystal violet, and photographed. Utilizing the ImageJ software, the quantification of both the percentage and intensity of crystal violet-stained cell colonies covering the area was conducted.

#### Cell Viability

LN229 cells were seeded at a density of 1.0 × 10^3 cells per well in a 96-well plate. Following seeding, the cells were treated with a specific concentration of CT for 72 hours. After the treatment period, MTT reagent was added to each well and incubated for 3 hours to allow the formation of Formazan crystals. Following incubation, DMSO was used to dissolve the Formazan crystals, and the absorbance was measured at 420 nm. The obtained absorbance values were used to generate a line graph representing the cell viability data.

#### Immunofluorescence staining of fixed cells

Cells were seeded on coverslips in 12-well plates and allowed to adhere. Upon completion of the experiment, the cells were fixed using 4% paraformaldehyde and permeabilized with PBS containing 0.25% Triton-X100. Subsequently, the cells were washed with PBS and blocked in a PBS solution containing 1% BSA, 0.3% Triton X-100, and 5% goat serum for 2 hours at room temperature. After blocking, primary antibodies diluted in the blocking buffer were added to the coverslips and incubated at 4°C overnight. Following primary antibody incubation, cells were washed three times with PBS for 5 minutes each. Fluorescence-conjugated secondary antibodies, prepared in blocking buffer, were added to the cells for 3 hours, followed by another round of washing with PBS. The cells were counterstained with DAPI (1 µg/ml) for 5 minutes at room temperature. Finally, coverslips were mounted onto glass slides using an anti-fade reagent as a mounting medium and imaged using a Zeiss LSM 880 confocal microscope.

#### Intracranial injection of GBM cells

Cells were collected using incomplete DMEM or F12 media, depending on the specific cell type intended for injection. Either 0.1 × 10^6 DBT-Luc GSCs or 3×10^5 U87-Luc cells, and 0.15×10^6 were intracranially injected into each animal’s hippocampus at a depth of 2.5 mm using a stereotaxic apparatus. Imaging of the animals commenced on the fifth day post-injection and continued every 5–6 days until the experiment’s conclusion. CT treatment began seven days post-injection, with animals receiving a daily intranasal dose of 4 IU CT (2 IU in the morning and 2 IU in the evening) every day for the next 25 days. Intranasal drug delivery was performed following the protocol mentioned earlier **[2]**.

#### In vivo Imaging

In vivo Bioluminescence imaging was conducted using the PerkinElmer IVIS Spectrum system under mild gas anaesthesia, employing isoflurane for the animals.

#### Haematoxylin and Eosin staining

Following perfusion, mouse brains were embedded in paraffin blocks and sectioned into 5 μm slices using the Leica Biosystems microtome instrument. These sections were affixed onto glass slides, deparaffinized, rehydrated, and subsequently stained with Harris Hematoxylin (Merck, 6159380051046) for nuclear visualization and Eosin Y (SDFCL, 44027G25) for cytoplasmic staining. After staining, the sections were mounted using DPX mounting medium and imaged at 0.8× magnification using a Lawrence and Mayo digital microscope.

#### Cy3 Labelling of Calcitonin Peptide

Labelling of Calcitonin Peptide is done according to the manufacturer’s protocol. Briefly, Cy3 NHS Ester (Cytiva PA13101) and Calcitonin Salmon (MedChemExpress HY-P0090) were incubated together overnight at 4°C with 1x PBS (pH=7.2) and 1mM of Sodium bicarbonate in a 5:1 mM ratio. After overnight incubation, the labelled peptide is purified through reverse-phase HPLC.

#### Intranasal delivery of labelled peptide

Tumour-bearing nude mice were administered PBS, unlabelled calcitonin (10 µg), and Cy3-labelled calcitonin (5 µg & 10 µg) via intranasal delivery. After 4 hours of administration, mice underwent whole-body fluorescence imaging with a 528nm excitation filter via the IVIS system. Immediately after whole body fluorescence imaging, all the mice were anaesthetised using isoflurane, and major organs (Brain, Lungs, Heart, kidneys, Liver, and Spleen) were harvested and subjected to organ-specific fluorescence imaging using the same excitation filter.

#### Sub-acute toxicity of calcitonin in C57BL/6 mice

C57BL/6 (6 weeks of age) were administered PBS or CT at doses of 1 I.U., 2 I.U., or 4 I.U. via intranasal delivery, morning and evening daily, for a consecutive 28 days. To eliminate sex differences, only female mice were used in the toxicity studies. Body weight, ECG recording, food consumption, and water consumption were measured every 7 days during the treatment. Blood samples and tissues were collected at 24 hrs following the final CT/PBS treatment. Blood samples for haematological analyses were collected in Microtainer Blood Collection Tubes with K2EDTA. Peripheral blood was analysed, and the haematological and biochemical parameters were analysed. Major organs, including the brain, lung, heart, liver, spleen, and kidney, were harvested from mice after the completion of CT/PBS treatment. Histological slides prepared from these organs were independently analysed by pathologists.

#### Survival Analysis

The survival analysis employing the Kaplan-Meier method was done using GraphPad Prism software (GraphPad Software, San Diego, CA).

#### Quantification and Statistical Analysis

Bar diagrams and Kaplan Meier plots were generated using GraphPad Software. Statistical analysis involved the calculation of p-values through unpaired t-tests with Welch’s correction or Student’s t-tests using Microsoft Excel, where indicated. ANOVA p-values were computed using GraphPad Prism 5 software. The significance of the Kaplan-Meier survival analysis was determined via the log-rank test. Pearson’s correlation coefficient was employed to calculate correlation coefficients using GraphPad Software. Experimental replicates, except those involving animal models, were conducted at least three times with duplicates for each experimental condition. Data are presented as mean ± standard deviation. A p-value <0.05 was considered significant, denoted by *, **, ***, and **** for p-values <0.05, 0.01, 0.001, and 0.0001, respectively, while “ns” indicates insignificant.

#### Molecular Modelling of CALCR

**Insilico reconstitution of full-length CALCR from structurally solved fragments**

For our simulation study, we reconstituted the full-length CALCR using the available structures of the transmembrane region of the CALCR (PDB ID: 6NIY) **[3]** and the solved CT-bound extracellular domain (PDB ID:5IIO **[4]**). Both of the above-mentioned PDBs are taken as the template, and the full-length CALCR was modeled using the MODELLER tool **[5]**. In the PDB: 5II0, the ECD of CALCR is bound to the unstructured region of the C-terminus of CT, while in another cryo-EM solved structure (PDB:6NIY), the TM region of CALCR is bound to the helical N-terminal region of CT. By carefully superimposing both structures onto our model, we were able to find the exact position and orientation of full-length functional CT when it’s bound to the full-length functional CALCR. While reconstructing the ECD for molecular simulations, we have included the disulfide bonds as per the available experimental information **[3]**. Also, we have deleted the first 24 residues that make up the protein’s signal peptide region. We have modeled the CALCR till residue number 428, as a recent study from the Patrick Sexton (Monash University, Australia) group exhibited that the CALCR was fully functional with the deletion of the C-terminal till residue 428 **[3]**. We chose the membrane composition to mimic the astrocyte membrane. The following composition was used: POPC:CHOL:SOPS:POPE:PSM:PIP2: 48%:20%:10%:10%:7%:5%.

#### Insilico reconstitution of mutant CALCR with binding partners

Along with the WT, we also modeled seven different mutant states of the protein. Three of these single point mutations (R45Q, A51T, P100L) are in the ECD domain, A307 is in TM5, V250M mutant is located on the ICL2, and the last two mutants (R404C, R420C) are in the amphipathic TM8. Besides constructing the seven mutant and WT systems for the CALCR that act as our reference for the mutant states, we have also constructed separate CALCR systems in complex with CT and G-proteins, both for the WT and mutant systems, based on the requirements of our study. For example, we have CT-bound ECD-mutant systems along with the CT-bound WT, where the WT CT-bound system acts as a control for changes observed in CT-binding due to mutations. Similarly, we have created WT CALCR and mutant V250M in complex with the G-proteins since ICL2, which carries the amino acid at the 250^th^ position, is in contact with the G-protein.

#### All-atom molecular dynamics simulation details

All the systems are built using the CHARMM-GUI input generator **[6–8]**. The CALCR was inserted in a membrane size of 15 nm x 15 nm with the above-mentioned lipid composition. The protein-bilayer system was then solvated with the TIP3P water model with roughly 22 Å water padding. The full system was charge-neutralised with 150 mM of Na and Cl ions.

The CHARMM36m forcefield was used as a particle definition and its interaction parameters **[9]**, and the GROMACS engine was used as the molecular simulation software **[10–12]**. The minimisation step was performed by the steepest descent algorithm, along with H-bond constrained using the LINCS algorithm. Then, two stages of NVT equilibration were performed with the Berendsen thermostat at a temperature of 303.15 K. Afterwards, four stages of NPT equilibration were done with the Berendsen barostat at 1 atm pressure. The further production run was extended for 600 ns to 1 μs with a time step of 2 fs for different systems of CALCR with Nose-Hoover thermostat at 303.15 K and Parrinello-Rahman barostat at 1 atm. All the analyses were carried out using the MD analysis modules **[13, 14],** and the molecular figures were rendered with the help of the ChimeraX tool **[15, 16].** For the binding energy calculation between CALCR and Gα, the gmx MMPBSA tool was used **[17, 18].**

All our data, including the simulated trajectories and input files for our AAMD simulations for all the systems under consideration, are publicly available on our GitHub repository: https://github.com/codesrivastavalab/GPCR-CalcitoninReceptor-AAMD. The large trajectories are deposited on a publicly available SharePoint location, and a link to the location is available in the above GitHub repository.

## Supplementary Figure Legends

**Supplementary Figure 1.**
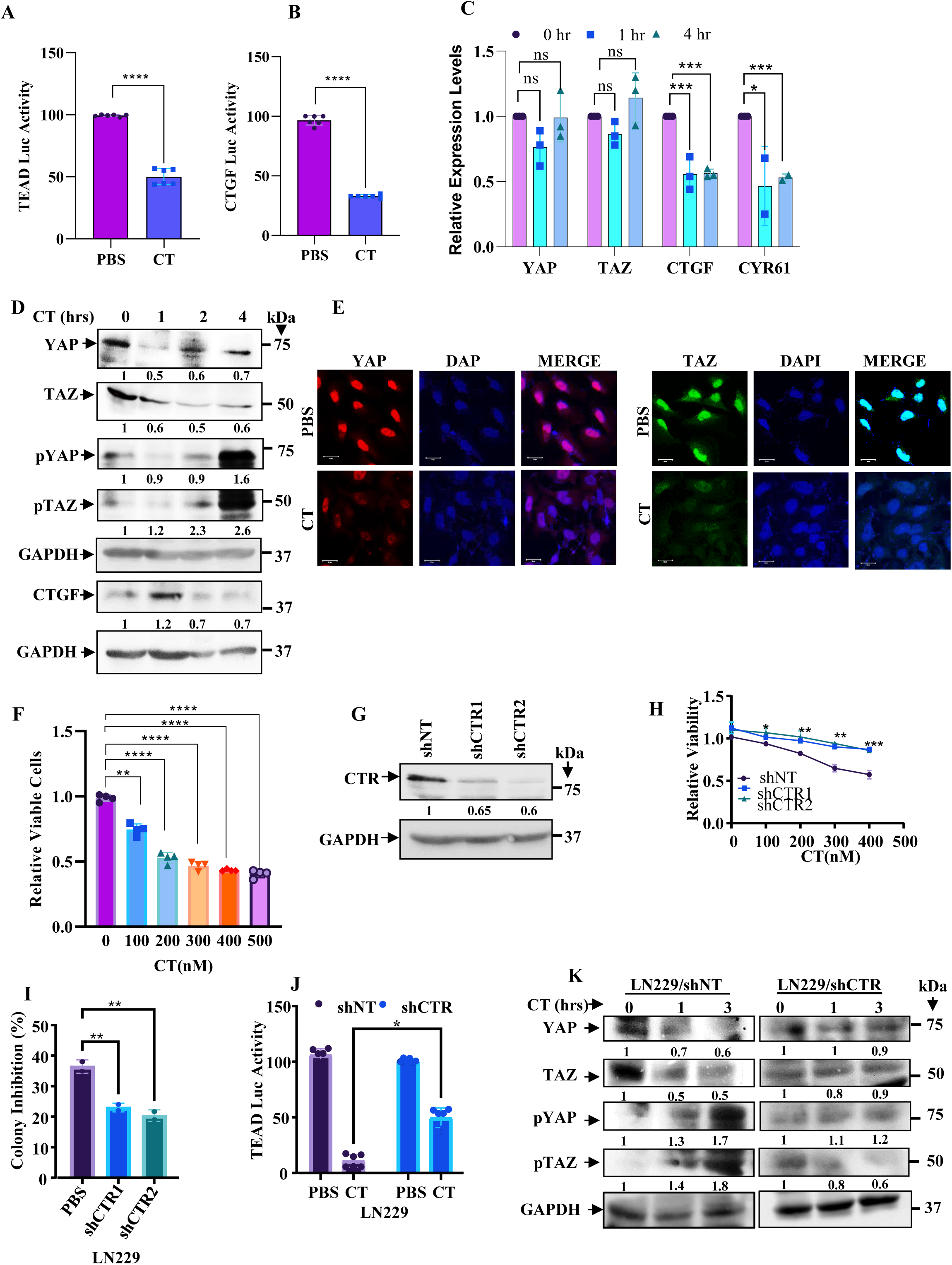
Inhibition of YAP and TAZ by CT in T98G cell line. Effect of PBS or CT on GBM cell line, T98G transfected with TEAD Luc (**A)** or CTGF Luc (**B)**. **(C)** RT-qPCR analysis shows YAP, TAZ, CTGF, and CYR61 transcript levels upon CT treatment (300nM) in the T98G cell line (two-way ANOVA; n=3/group). (**D)** Western blotting shows levels of YAP, TAZ, pYAP, pTAZ, and CTGF upon CT treatment in the T98G cell line. (**E)** Confocal microscopy analysis shows YAP and TAZ levels in T98G cells with and without CT treatment. Red indicates YAP, green indicates TAZ, and blue indicates DAPI. The merged images are shown for representation. Magnification x63, scale bar 20 μm. (**F)** MTT assay showing the viability of T98G cells at the indicated concentrations of CT after 72 hours of treatment (one-way ANOVA; n=4/group). **(G)** Western blot showing the knockdown of CTR in LN229 cells using two shRNAs. (**H)** MTT assay showing the effect of CT on LN229/ shNT and LN229/ shCTR cells at the indicated concentrations of CT (two-way ANOVA; n=4/group); n=3/group). (**I)** Quantification of growth inhibition by CT in shNT/LN229 and shCTR/LN229 conditions as seen in the colony formation assay (t-test; n=2/group). (**J)** TEAD Luc activity inhibition in shNT and shCTR knockdown conditions upon CT treatment (t test; n=6/group). (**K)** Western blotting shows YAP, TAZ, pYAP, and pTAZ levels upon CT treatment under shNT and shCTR conditions.

**Supplementary Figure 2.**
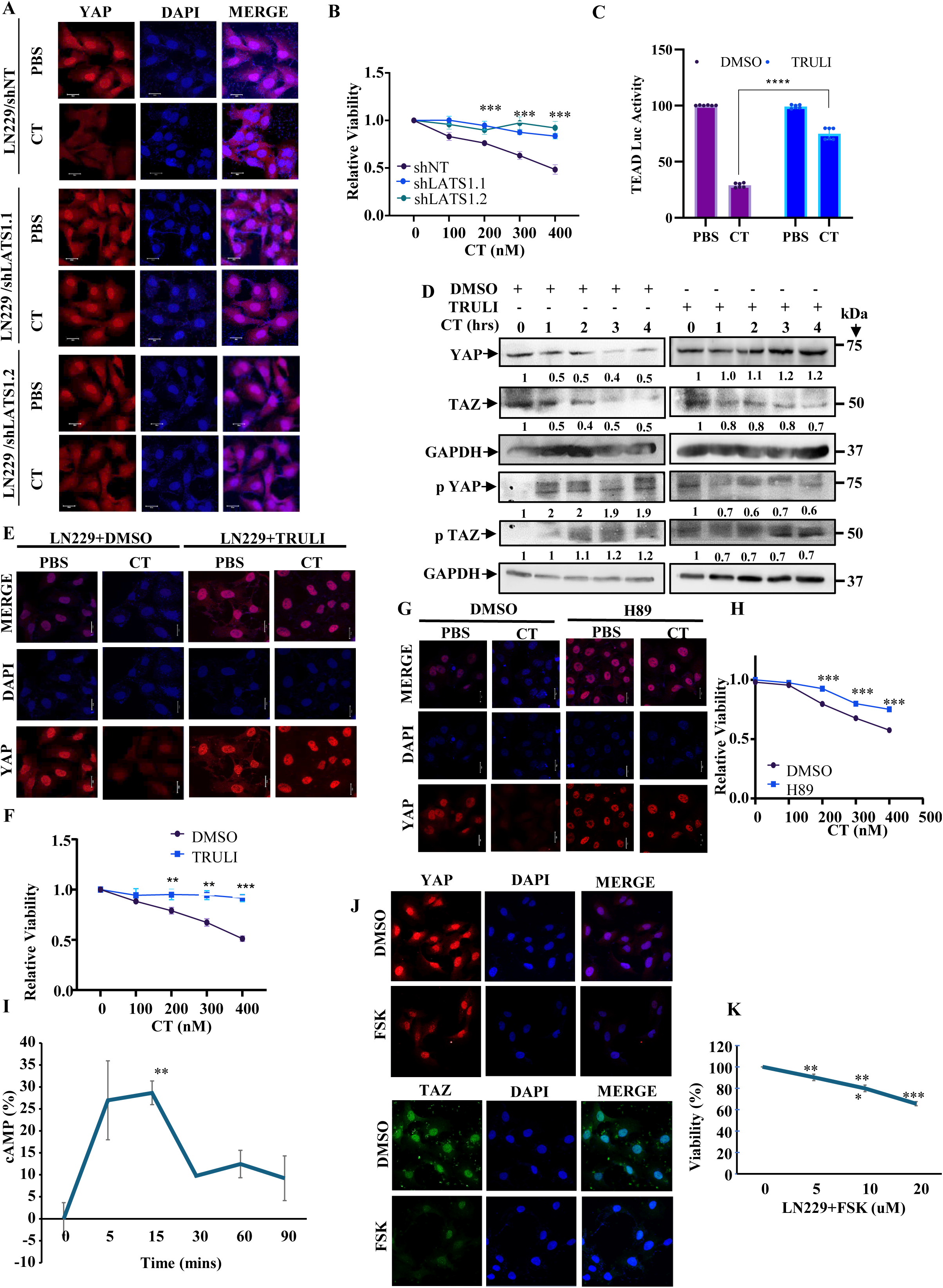
CT modulates YAP and TAZ in a cAMP-PKA-LATS1-dependent manner. (**A)** Confocal microscopy analysis shows YAP and TAZ levels in shNT and shLATS1 conditions with and without CT treatment. Red indicates YAP, green indicates TAZ, and blue indicates DAPI. The merged images are shown for representation. Magnification x63, scale bar 20 µm. (**B)** MTT Assay revealing Cell Viability in shNT and shLATS1 Cells following CT treatment at the indicated concentrations (two-way ANOVA; n=4/group). (**C)** LN229 cells were first transfected with the TEAD Luc construct and pre-treated with TRULI (10 µM) for six hours, followed by CT treatment for four hours before cell harvesting for luciferase activity measurement (two-way ANOVA; n=6/group). (**D)** LN229 cells were pre-treated with TRULI for six hours and treated with CT for the indicated time points. The lysates were assessed for total and phosphorylated YAP and TAZ. (**E)** Confocal imaging displays YAP and TAZ levels after CT treatment, with either DMSO or TRULI pre-treatment. (**F)** Cell viability of LN229 cells was assessed by MTT assay upon CT treatment with and without TRULI treatment (two-way ANOVA; n=3/group). (**G)** Confocal imaging shows YAP’s abundance and nuclear localisation upon CT treatment in DMSO or H89-pre-treated LN229 cells. (**H)** Cell viability of LN229 cells was assessed by MTT assay upon CT treatment in DMSO or H89-pretreated LN229 cells (two-way ANOVA; n=4/group). (**I)** Levels of cAMP abundance at the indicated time points after CT treatment were determined using the ELISA assay kit (t test; n=2/group). (**J)** Confocal images depicting the expression levels of YAP and TAZ upon FSK treatment. Red indicates YAP, green indicates TAZ, and blue indicates DAPI. The merged images are shown for representation. Magnification x 63, scale bar=20 µm. (**K)** Cell viability of LN229 cells with the indicated concentration of FSK treatment after 72 hours of treatment (t test; n=4/group).

**Supplementary Figure 3.**
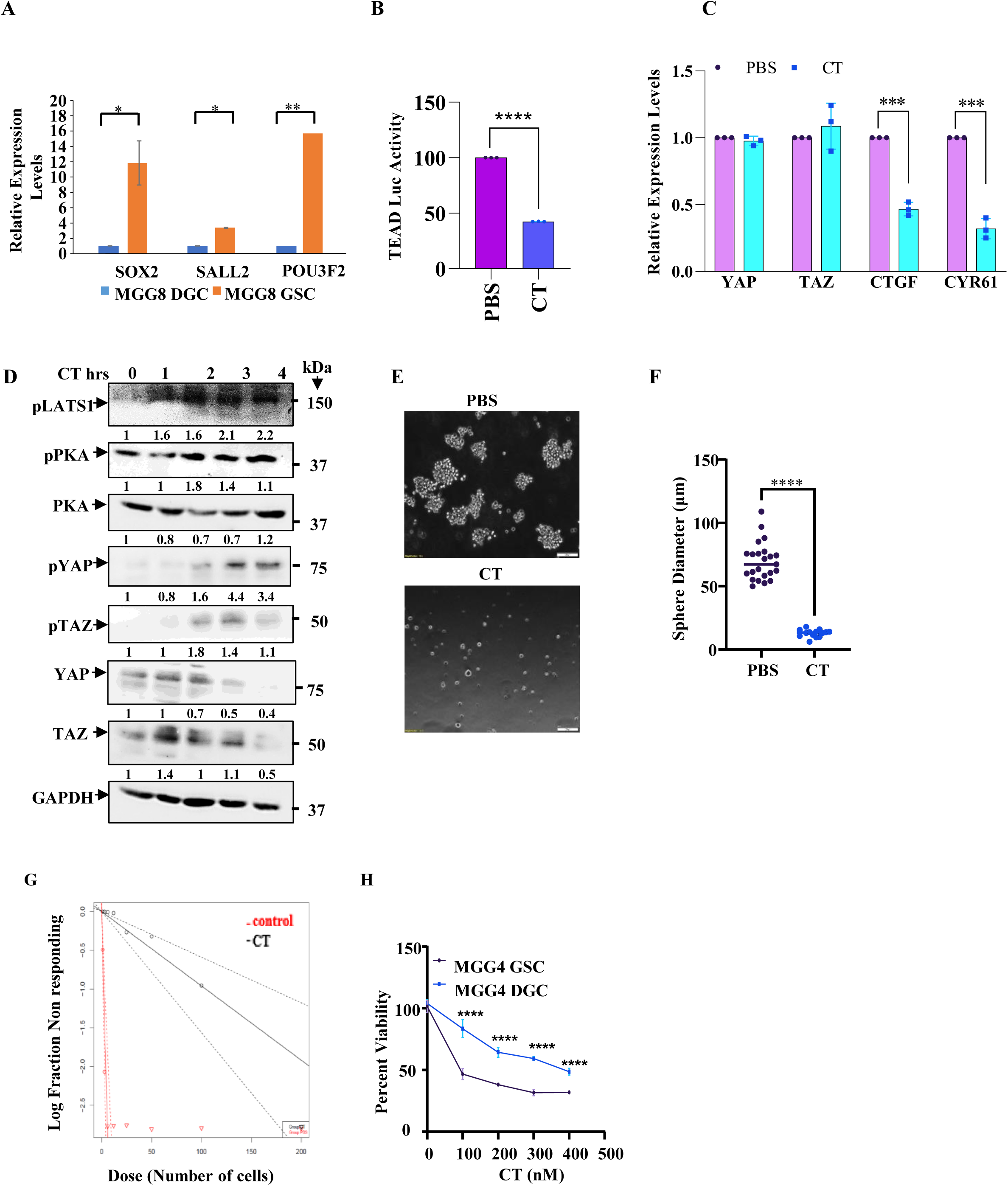
GSCs are better suited for CT-mediated growth inhibition. **(A)**. qRT-PCR confirms the GSC and DGC status based on the levels of reprogramming factors SOX2, SALL2, and POU3F2 (t test; n=2/group). **(B)** TEAD-Luc activity in GB-3 neurospheres treated with PBS or CT (t test; n=3/group). **(C)** mRNA expression levels of YAP, TAZ, CTGF, and CYR61 in GB3 cells treated with CT (n=3/group). **(D)** Western blotting shows the levels of YAP, TAZ, pYAP, pTAZ, pLATS1, pPKA, and PKA in GB3 spheres treated with CT at the indicated time points. **E, F,G** shows the impact of CT upon the growth of GB3 neurospheres by sphere formation assay and limiting dilution assay, respectively. **(H)** MTT assay shows the differential growth inhibition of CT on MGG4 GSCs vs DGCs (two-way ANOVA; n=4/group).

**Supplementary Figure 4.**
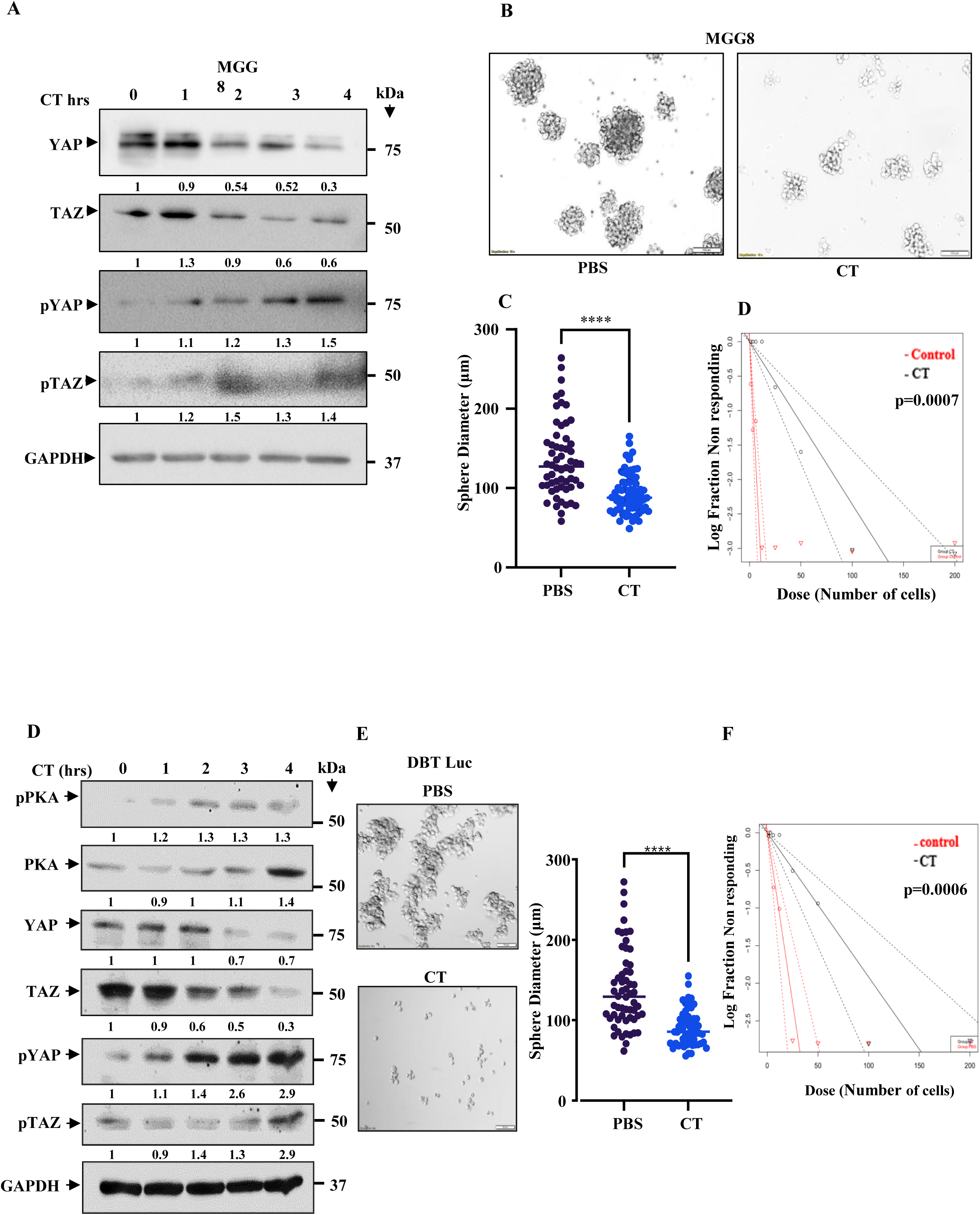
CT inhibits the growth of human and murine GSCs. **(A)** Western blot shows the levels of YAP, TAZ, pYAP and pTAZ upon CT treatment at the indicated time-points. Impact of CT on human-derived GSCs, MGG8, as shown by sphere formation assay **(B)** and limiting dilution Assay **(C). (D)** Western blotting shows the levels of YAP, TAZ, pYAP, pTAZ, PKA, and pPKA in murine GSCs, DBT-Luc. Impact of CT on human-derived GSCs, MGG8, as shown by sphere formation assay **(E)** and limiting dilution Assay **(F)**.

**Supplementary Figure 5.**
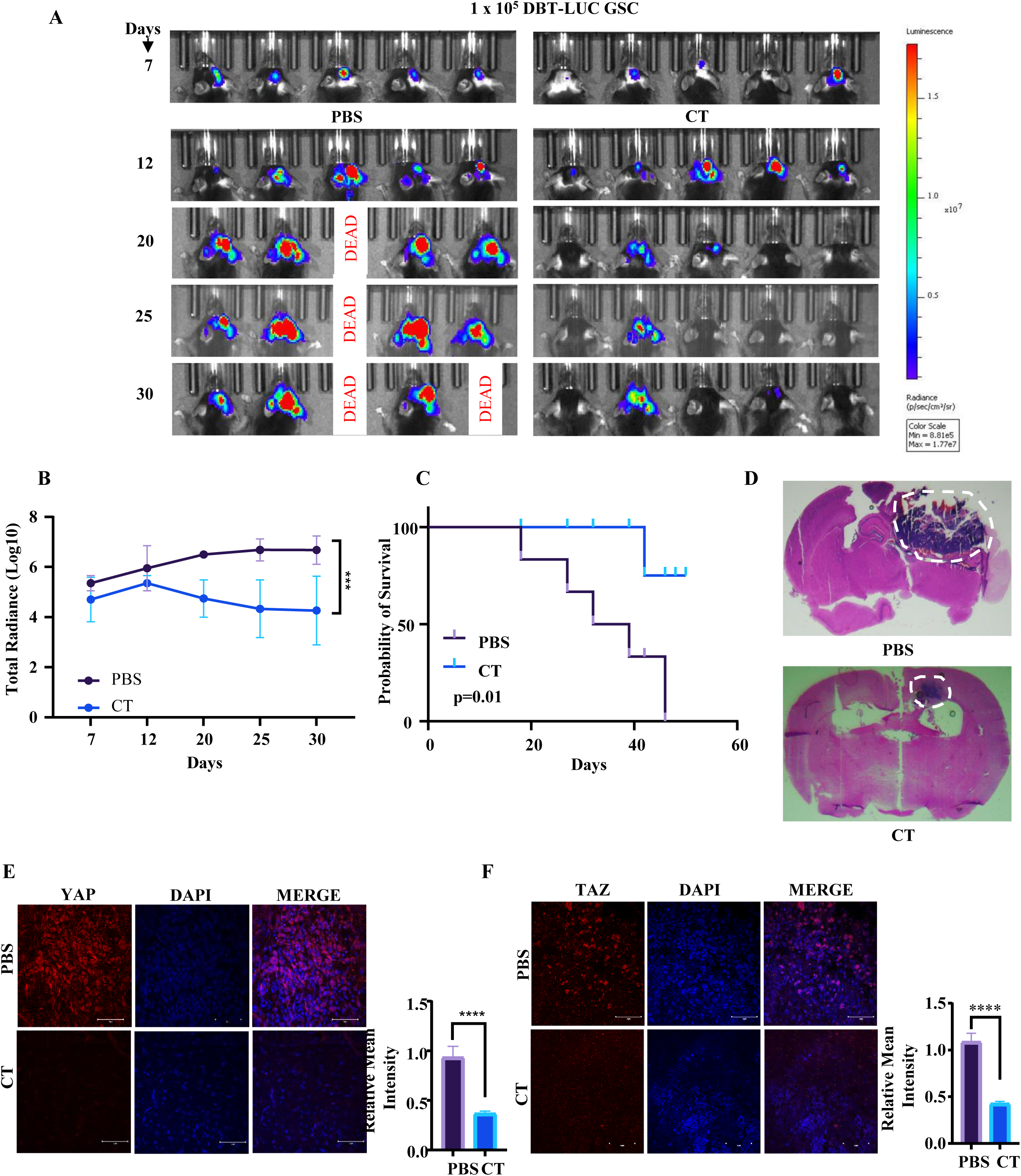
Intranasal CT administration inhibits DBT-Luc GSC-derived tumor in C57BL/6 mice. **(A)** 0.1×10^6^ DBT-Luc GSCs were injected intracranially into the brain of C57BL/6 mice (n=5 per group). The tumor was allowed to grow for seven days, after which CT was administered intranasally at a dose of 2 IU/ animal. In vivo, bioluminescence imaging of the animals was taken every five days from the date of injection. (**B)** The average radiance efficiency between the PBS and CT-administered groups is plotted (two-way ANOVA; n=5/group). (**C)** The Kaplan-Meier graph shows the survival difference between the two groups of mice that were administered PBS or CT (log rank test; n=5/group). (**D)** Haematoxylin and eosin staining shows a larger tumor (depicted by dark blue colour) in animals administered with intranasal PBS compared to a smaller tumor in the CT-administered group. Confocal microscopy shows YAP **(E)** and TAZ **(F)** levels in the PBS and CT-treated groups.

**Supplementary Figure 6.**
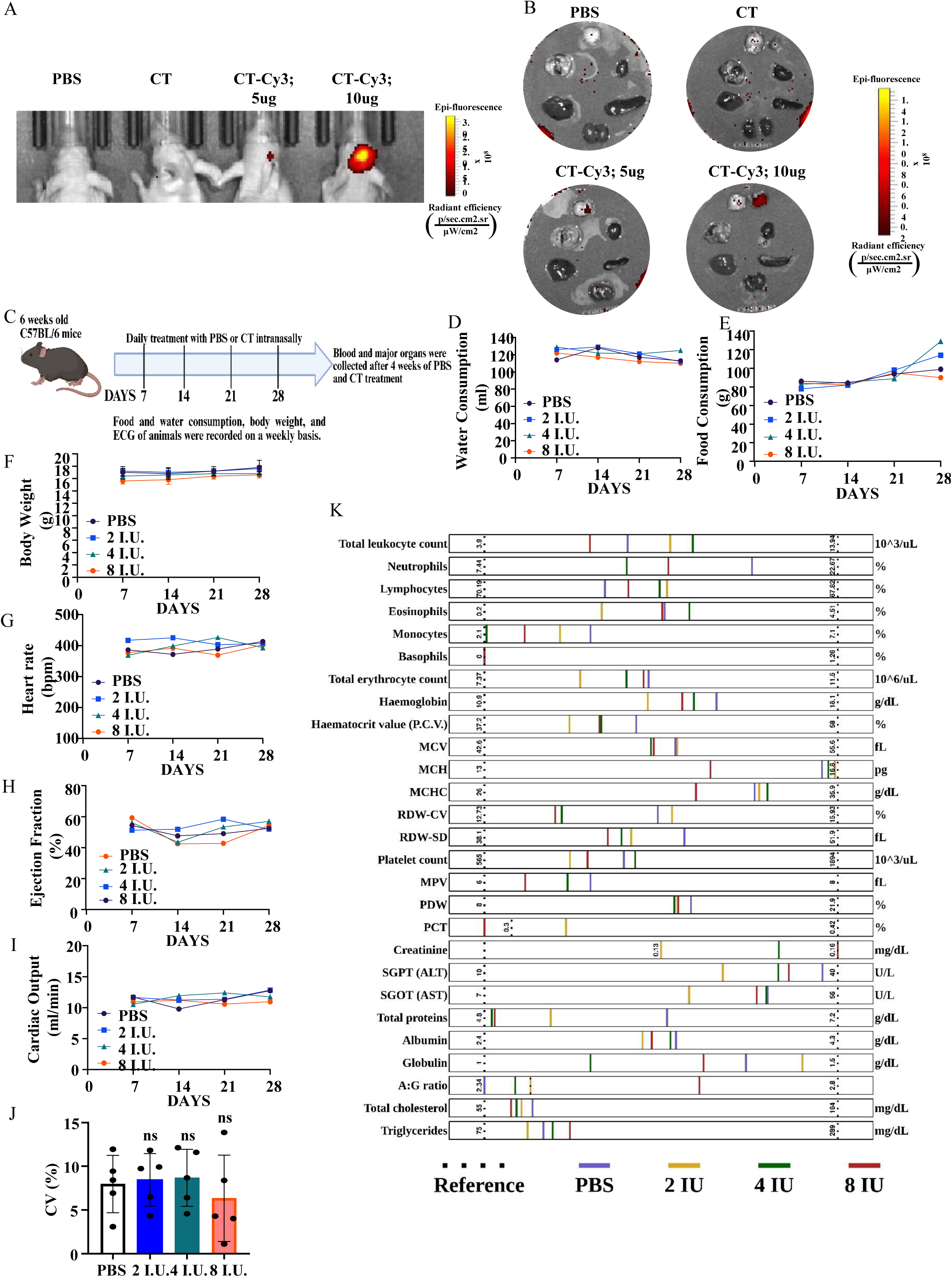
Bioavailability of CT in the brain upon intranasal administration and toxicity profile of CT in animals. **(A)** In vivo imaging shows fluorescence activity in the brain of animals after 3 hours of intranasal Cy3-tagged CT-treatment. **(B)** Fluorescence activity in the major organs of the animals. **(C)** The illustration depicts the strategy to study the sub-acute toxicity of CT in animals. Measures of **(D),** water consumption **(E)**, Food consumption **(F),** body weight, **(G)** heartbeat (beat per minute) **(H)**, ejection fraction (%) **(I)**, cardiac output (ml/min) **(J)** co-variance of heart rate **(K),** Haematological parameters of the animals measured 28 days post CT administration. For panels D, E, F, G, H, and I, one-way ANOVA; n=5/group.

**Supplementary Figure 7.**
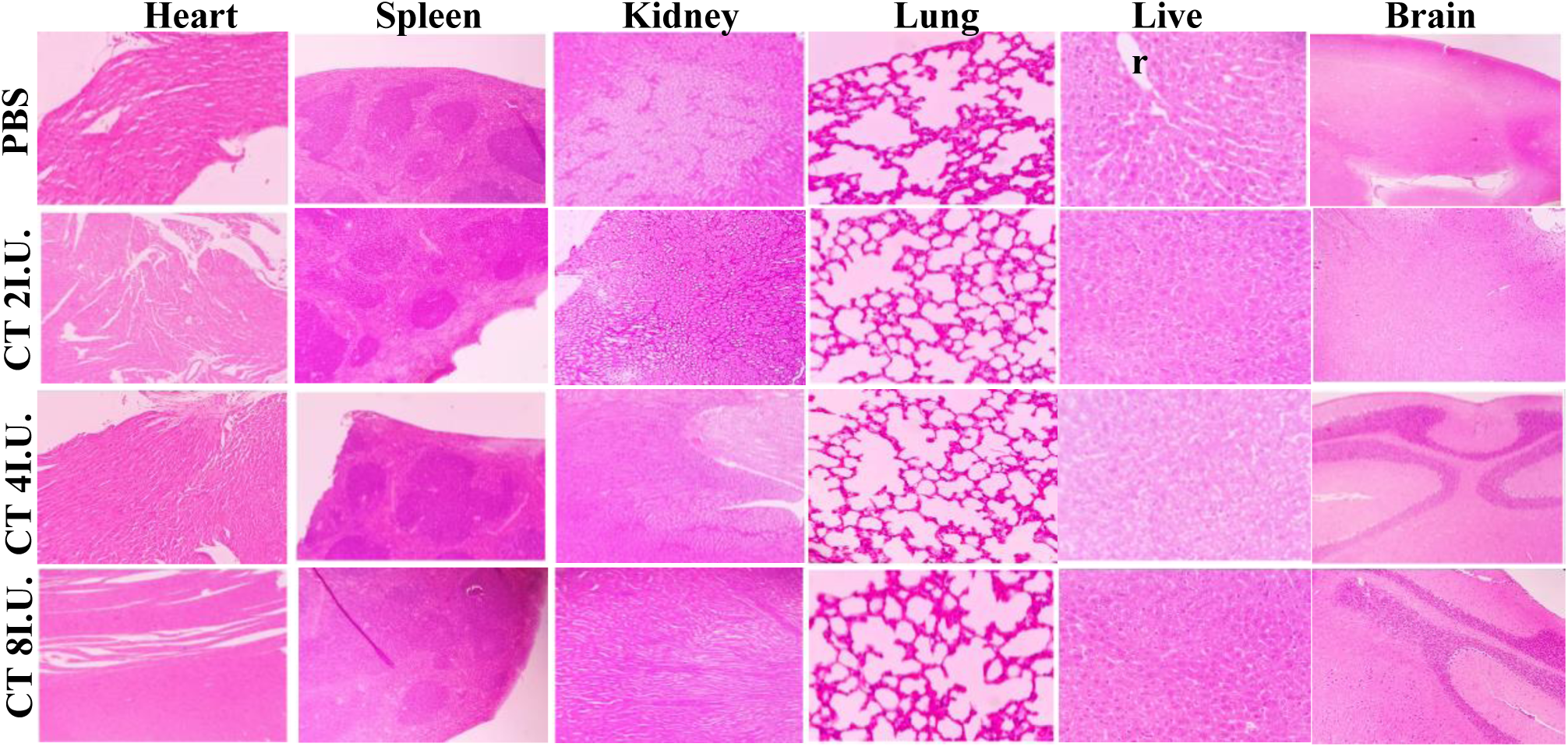
H&E staining of the major organs Heart, Spleen, Kidney, Lung, Liver, and Brain from C57BL/6 mice after 28 days of intranasal PBS or CT administration

**Supplementary Figure 8.**
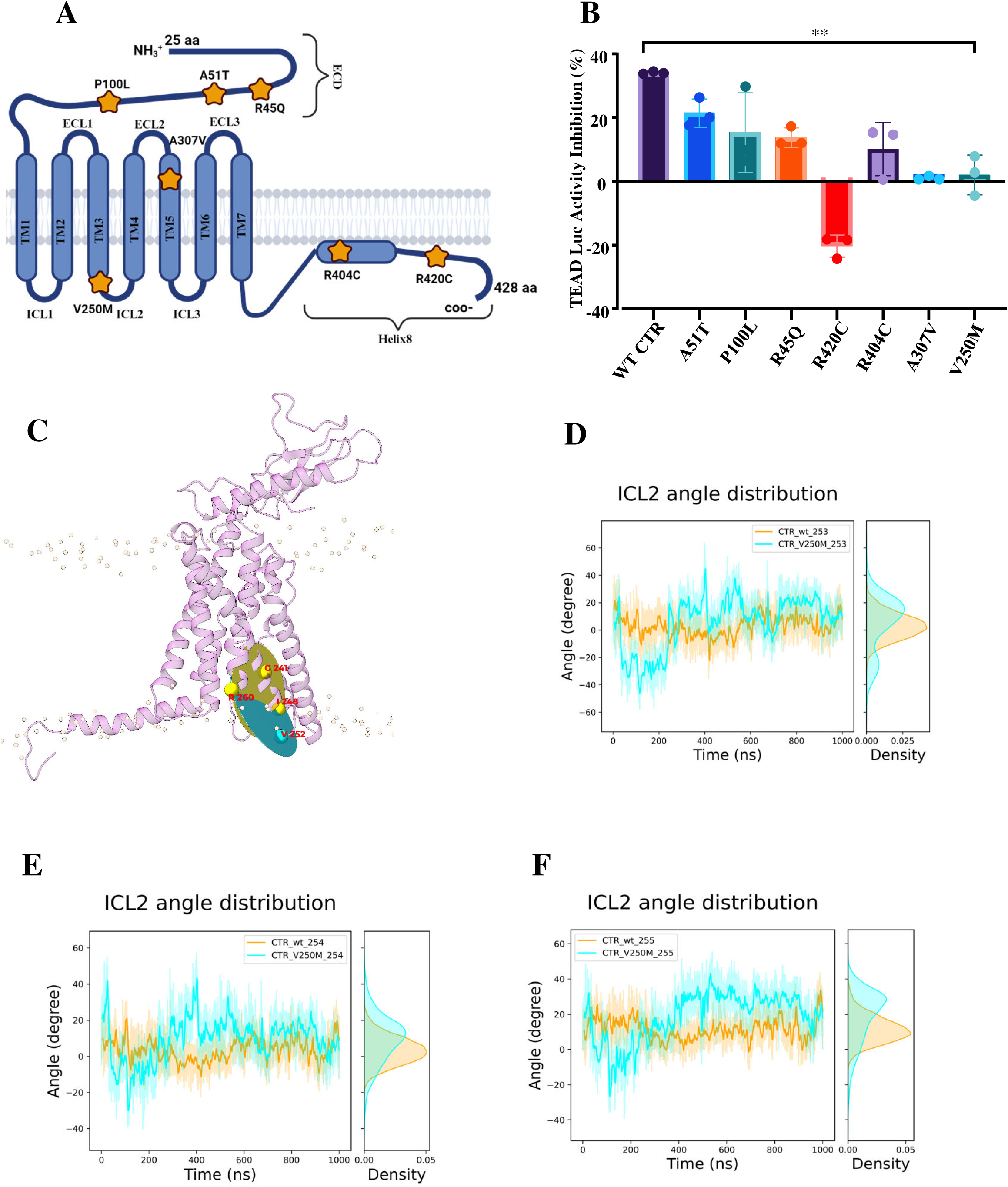
CALCR mutants lose the ability to activate the Hippo pathway, and the impact of the V250 M mutation on ICL2 flexibility. (A) The figure shows the cartoon representation of CALCR and the location of seven mutations denoted by yellow stars. (B) The graph shows the percentage inhibition of TEAD-Luc activity for the CALCR mutants compared to WT CALCR upon CT treatment. (C)The figure shows the two planes, i.e., plane-0 (yellow), defined by the c-alpha carbon of residues R260, I248, and G241, and plane-1 (cyan), defined by the c-alpha carbon of residues R260, I248, and V252. The plots show ICL2 angle distribution in the WT and V250M system with respect to residue (D) F253, (E) T254, and (F) E255.

**Supplementary Figure 9.**
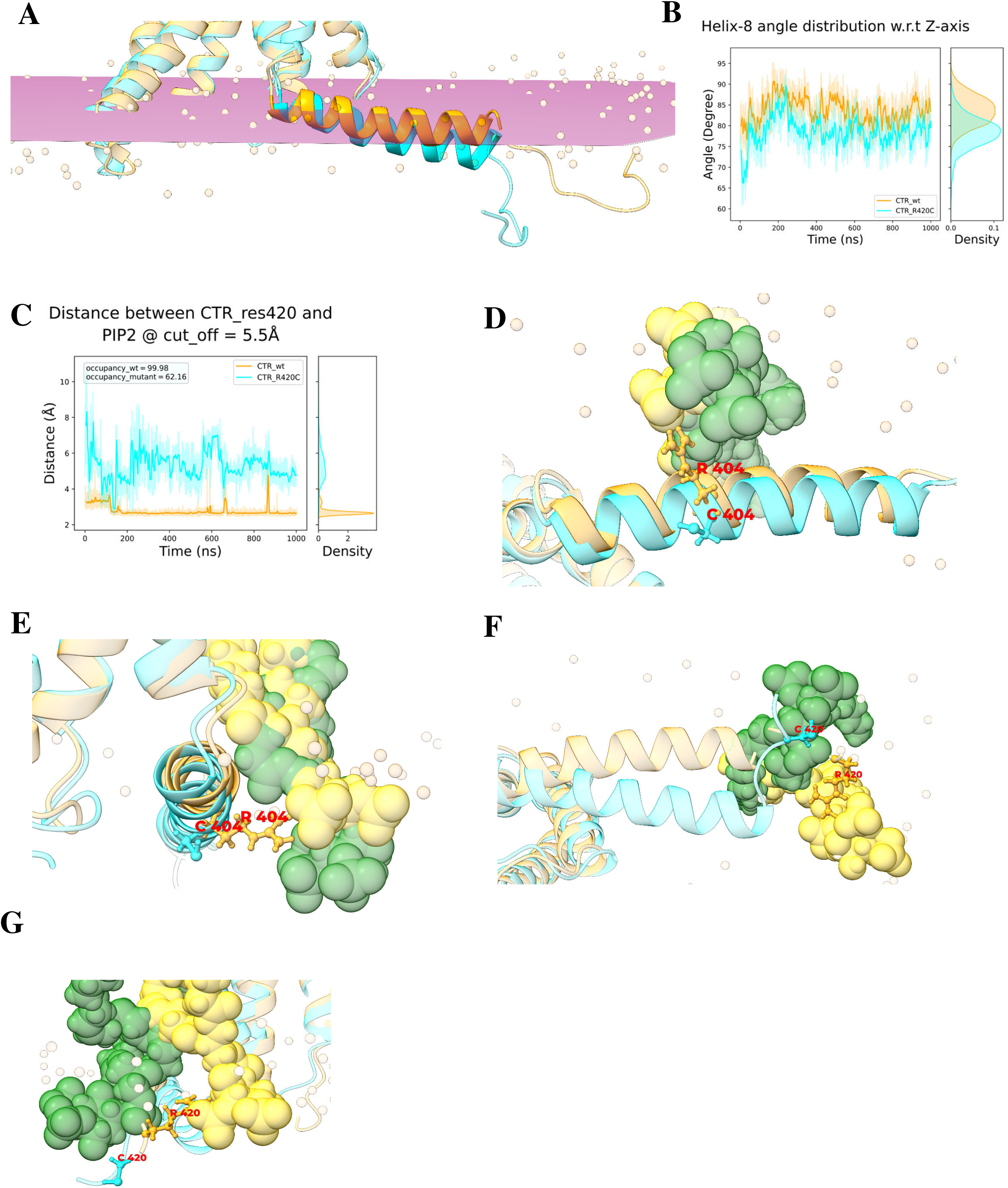
Illustration of R404C and R420C mutant orientation with respect to membrane PIP2 interaction and effect on the conformational shift and interaction with membrane PIP2 due to R420C mutation. (A) The superimposed image shows the orientation of the 8th helix in the WT CALCR (orange) and R420C mutant (cyan). (B) The plot shows the difference in the angle distribution of the 8th helix with the z-axis in the WT CALCR and R420C mutant. (C) The plot shows the interaction of the 420th residue with PIP2 lipids of the membrane in the WT CALCR and R420C mutant. (D) The figure shows the bottom view of residue 404 interacting with PIP2 lipid in WT CALCR (R404 in orange and PIP2 lipid in gold) and mutant CALCR (C404 in cyan and PIP2 lipid in green). (E) The figure shows a side view of the exact system mentioned in Figure D. (F) The figure shows the bottom view of the interaction of residue 420 with PIP2 lipid in both WT CALCR (R420 in orange and PIP2 in gold) and mutant CALCR (C420 in cyan and PIP2 in green). (G) The figure shows the side view of the exact system mentioned in Figure F.

**Supplementary Figure 10.**
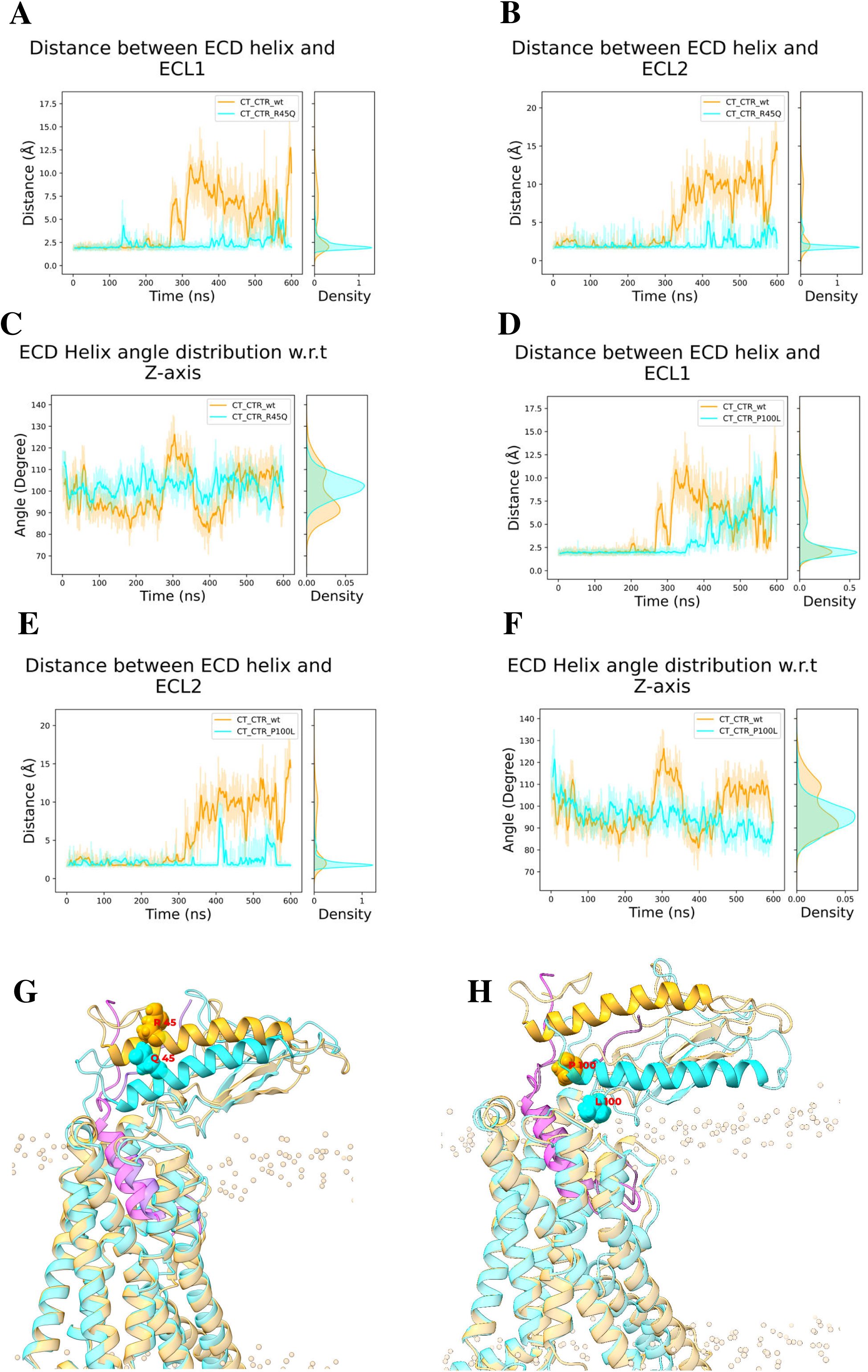
Illustration and quantification of ECD conformational shift due to the R45Q and P100L mutation. **(A)** The plot shows the minimum distance between the ECD helix and ECL1 loop for WT CALCR and R45Q mutant when bound to CT. **(B)** This plot shows the minimum distance between the ECD helix and ECL2 loop for the previously mentioned systems. **(C)** The plot shows the ECD helix angle with respect to the Z-axis for the systems in the same way as Figure A. **(D)** The plot shows the minimum distance between the ECD helix and ECL1 loop for WT CALCR and P100L mutant when bound to CT. **(E)** This plot shows the minimum distance between the ECD helix and ECL2 loop for the previously mentioned systems. **(F)** The plot shows the ECD helix angle with respect to the Z-axis for the systems, the same as Figure D. **(G)** The superimposed image shows the ECD orientation between WT CALCR (orange) and R45Q mutant (cyan) when bound to CT (pink). **(H)** The superimposed image shows the ECD orientation between WT CALCR (orange) and P100L mutant (cyan) when bound to CT (pink).

**Supplementary Figure 11.**
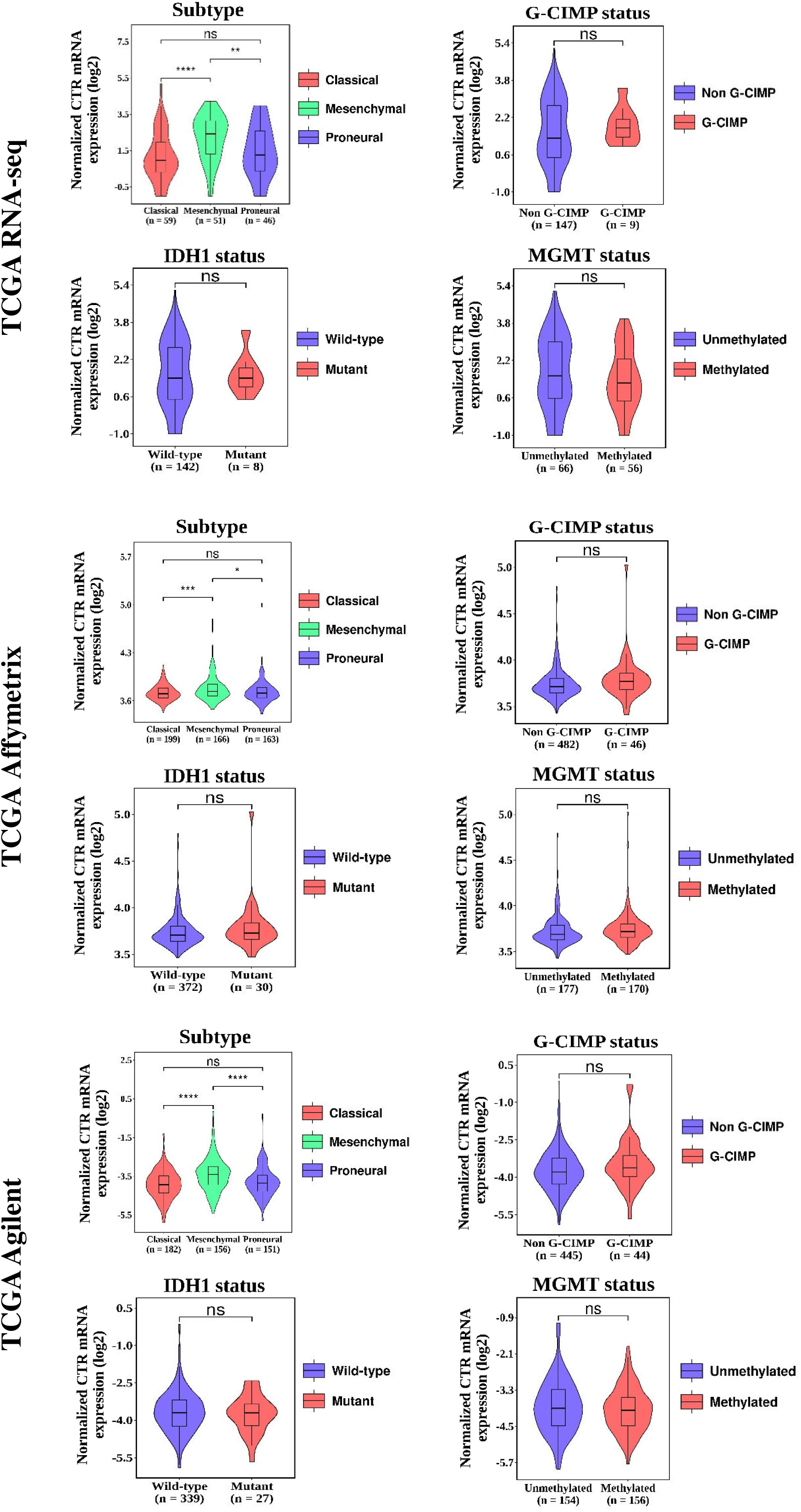
CALCR Expression Levels in different TCGA Datasets. Expression levels of *CALCR* are shown across different TCGA datasets. The analysis includes comparisons based on molecular subtypes, G-CIMP status, IDH mutation status, and MGMT promoter methylation.

